# Oral vancomycin treatment alters levels of indole derivatives and secondary bile acids modulating the expression of mTOR pathway genes in astrocytes during EAE

**DOI:** 10.1101/2024.06.14.599110

**Authors:** Paola Bianchimano, Paola Leone, Emma M. Smith, Cristina Gutierrez-Vazquez, Erli Wind-andersen, Gerold Bongers, Sebastian Cristancho, Howard L. Weiner, Jose C. Clemente, Stephanie K. Tankou

## Abstract

Astrocytes play important roles in the central nervous system (CNS) during health and disease. Prior studies have shown that gut commensals derived indole derivatives as well as secondary bile acids modulate astrocyte function during the late stage of EAE (recovery phase). Here we show that administering vancomycin to mice starting during the early stage of EAE improved disease recovery, an effect that is mediated by the gut microbiota. We observed that 6 taxa within the *Clostridia vadin BB60* group were enriched in vancomycin treated mice compared to untreated EAE mice. Vancomycin-treated EAE mice also had elevated serum levels of the anti-inflammatory tryptophan derived metabolite, indole-3-lactic acid and decreased levels of deoxycholic acid, a pro-inflammatory secondary bile acid. RNA sequencing revealed altered expression of several genes belonging to the mammalian target of rapamycin (mTOR) pathway in astrocytes obtained during the late stage of EAE from vancomycin treated EAE mice. Furthermore, we observed a link between serum levels of indole derivatives and bile acids and expression of several genes belonging to the mTOR pathway. Interestingly, the mTOR signaling cascades have been implicated in several key biological processes including innate (e.g., astrocyte) immune responses as well as neuronal toxicity/degeneration. In addition, rapamycin, a specific inhibitor of mTOR, has been shown to inhibit the induction and progression of established EAE. Collectively, our findings suggest that the neuroprotective effect of vancomycin is at least partially mediated by indole derivatives and secondary bile acids modulating the expression of mTOR pathway genes in astrocytes.

**HIGHLIGHTS:**

- Vancomycin attenuated established EAE through regulation of the microbiota
- Vancomycin induced increased serum levels of indole-3-lactic acid, decreased serum levels of indoxyl-3-sulfate, p-cresol and decreased stool levels of deoxycholic acid
- Vancomycin modulated the expression of mTOR pathway genes in astrocytes
- *Lactobacillus reuteri* regulated the expression of mTOR pathway genes in astrocytes
- Serum levels of indole-3-lactic acid, indoxyl-3-sulfate, p-cresol and deoxycholic acid correlated with expression of mTOR pathway genes in astrocytes

## INTRODUCTION

Multiple sclerosis (MS) is a chronic central nervous system (CNS) inflammatory and neurodegenerative disease and one of the leading causes of disability in young individuals. MS is typically diagnosed between the ages of 20-40 as relapsing-remitting MS and 15-20 years later, some of these patients are going to enter the progressive phase of the disease which is non-remitting. Several drugs are FDA approved for the treatment of MS and are effective at suppressing relapsing disease but have limited impact on disease progression. During the progressive phase of MS, the inflammatory response is confined to the CNS and is primarily driven by astrocytes and microglia^1,2^. Astrocytes are the most abundant cell population in the CNS, and they have several functions including modulation of the blood-brain barrier, regulation of neuronal transmission and CNS development and repair^3–6^. Astrocytes have also been implicated in the pathogenesis of several CNS inflammatory and neurodegenerative diseases including MS and its mouse model^7–10^. One possible reason why the current immunomodulatory therapies failed in progressive MS is because they do not effectively target astrocytes or microglia. Hence, developing novel disease modifying therapeutic strategies is crucial to slowing cognitive decline and enhancing the quality of life of progressive MS patients.

The gut microbiome plays an important role in autoimmunity including MS and its mouse model, experimental autoimmune encephalomyelitis (EAE)^11–14^. One study reported that dietary tryptophan is metabolized by the gut microbiota into indole and its derivatives which are aryl hydrocarbon receptor (AhR) agonists acting on astrocytes to limit CNS inflammation and neurodegeneration in EAE mice^15^. The same study found that oral administration of ampicillin but not vancomycin during the late stage of EAE interfered with disease recovery by modulating astrocyte function via alteration in gut derived AhR agonist levels, namely indole-3-propionic acid, indoxyl-3-sulfate and indole-3-aldehyde^15^. Hence, indole derivatives are important regulators of the gut-brain axis during EAE. While human MS differs in many ways from EAE, multiple MS-associated microorganisms exacerbate the EAE phenotype and EAE mice colonized with MS derived microbiota develop more severe disease than EAE mice reconstituted with microbiota derived from healthy donors^16,17^. These findings demonstrate that aspects of EAE reflect clinically relevant host-microbiome interactions, providing mechanistic insights essential to interpret the large-scale studies correlating aspects of the gut microbiome with MS in patients.

The identity of gut commensals modulating astrocyte function remains obscure. We have previously shown that oral administration of vancomycin prior to EAE induction ameliorates disease and demonstrated that this protective effect is mediated via the microbiota^13^. In the present study, we showed that EAE mice fed microbiota derived from vancomycin treated mice post-symptom onset had better recovery than mice fed microbiota from untreated mice. We observed that administration of vancomycin post-EAE induction but before symptoms onset (early stage of EAE) improved disease recovery. Microbiome characterization sequencing revealed enrichment of the genus *Clostridia vadin BB60 group* in vancomycin treated EAE mice. Vancomycin and ampicillin treated mice had decreased serum levels of indoxyl-3-sulfate (3-IS), indole-3-propionic acid (3-IPA) and p-cresol compared to untreated EAE mice. Vancomycin but not ampicillin treated EAE mice had increased serum indole-3-lactic acid (3-ILA) levels compared to untreated EAE mice. Interestingly, we observed a positive correlation between serum levels of ILA and the abundance of 4 taxa belonging to the *Clostridia vadin BB60 group* genus. We also found decreased serum levels of most secondary bile acids including deoxycholic acid (DCA), a pro-inflammatory bile acid, in vancomycin treated EAE mice. Serum levels of DCA positively correlated with the abundance of 6 taxa belonging to the *Muribaculaceae* genus which was enriched in untreated EAE mice compared to vancomycin treated EAE mice.

Given that indole and its derivatives are AhR agonists that have been shown to modulate astrocyte function during EAE, we conducted RNA sequencing on astrocytes derived from these EAE mice. RNA sequencing of astrocytes revealed that vancomycin altered the expression of multiple genes including genes belonging to the mTOR pathway. We observed a correlation between levels of expression of genes belonging to the mTOR pathway in astrocytes and serum levels of indole derivatives and bile acids. Furthermore, we have previously observed increased abundance of *Lactobacillus reuteri*, an indole-3-lactic acid producing bacteria, in vancomycin treated mice. In the current study, we found increased abundance of *Lactobacillus reuteri* in vancomycin treated naïve mice. We showed that mice supplemented with *L. reuteri* prior to EAE induction or at the time of symptoms onset had better disease recovery compared to EAE mice who received vehicle. We also performed RNA sequencing from astrocytes isolated from mice receiving *L. reuteri* or vehicle. We observed alteration in the expression of genes belonging to the mTOR pathway including increased expression of Acsl3 in mice supplemented with *L. reuteri* which was also increased in vancomycin treated EAE mice. Interestingly, rapamycin, a specific inhibitor of the mTOR pathway ameliorates EAE by inducing T regulatory cells or suppressing Th1 and/or Th17 cells. However, the role of the mTOR pathway in astrocytes during EAE remains poorly understood.

Our results suggest that mTOR pathway genes in astrocytes could be potential targets for therapeutic manipulation and provide a foundation to explore the mechanisms by which gut commensals derived metabolites modulate this pathway in astrocytes during chronic CNS inflammation.

## MATERIALS AND METHODS

### Mice

C57BL/6J female mice of 8-10 weeks of age were purchased from the Jackson Laboratory and housed in a specific pathogen-free facility at the Icahn School of Medicine at Mount Sinai or the Harvard Institute of Medicine. Animals were kept on a 12-h light/dark cycle with access to food (PicoLab Rodent Diet 5053) and distilled water at neutral pH provided by the animal facility.

Mice were co-housed, 3-5 mice per cage from the same experimental condition. Animals were allowed to acclimate for a total of 5 days before initiation of treatment. Upon arrival at the animal facility, mice were housed in groups of 3-5 per cage by the mouse facility staff. Two days after arrival, mice are re-grouped so that each experimental group receives mice from all the cages that were set up on arrival. At this age, mice have not undergone any experimental manipulations and are indistinguishable from one another. Groups are formed by redistributing the mice and exchanging bedding.

Animals were housed in a biosafety level 2 facility using autoclaved cages and aseptic handling procedures. All animal experiments were approved by the Institutional Animal Care and Use Committee (IACUC) at Icahn School of Medicine at Mount Sinai or Harvard Medical School and carried out in accordance with those approved animal experiment guidelines.

### EAE induction

EAE was induced in 8- to 10-week-old female C57BL/6J mice by injecting 250 μg of MOG_35-55_ peptide (Genemed Synthesis) emulsified in complete Freund’s adjuvant (CFA) (BD Difco) per mouse, injected subcutaneously into each flank, followed by intraperitoneal administration of 150 ng of pertussis toxin (List Biological Laboratories, Inc.) per mouse on day 0. A second dose of 300 ng pertussis toxin was administered on day 2. Clinical signs of EAE were assessed according to the following scale: 0, no signs of disease; 1, loss of tone in the tail; 2, hind limb paresis; 3, hind limb paralysis; 4, tetraplegia; and 5, moribund. Differences between the groups were determined by Friedman test and Dunn’s correction for multiple comparisons.

### Antibiotic treatments

For experiments assessing the impact of the time of initiation of vancomycin treatment on EAE development (Figure 1A), three treatment groups were subjected to different time courses of vancomycin administration: −14 to 0 days post immunization (DPI) (vanco prior), 10 to 30 DPI (vanco post), −14 to 30 DPI (vanco continuous). Vancomycin 0.5mg/mL (Research Products International Corp.) was administered in drinking water.

**Figure 1.**
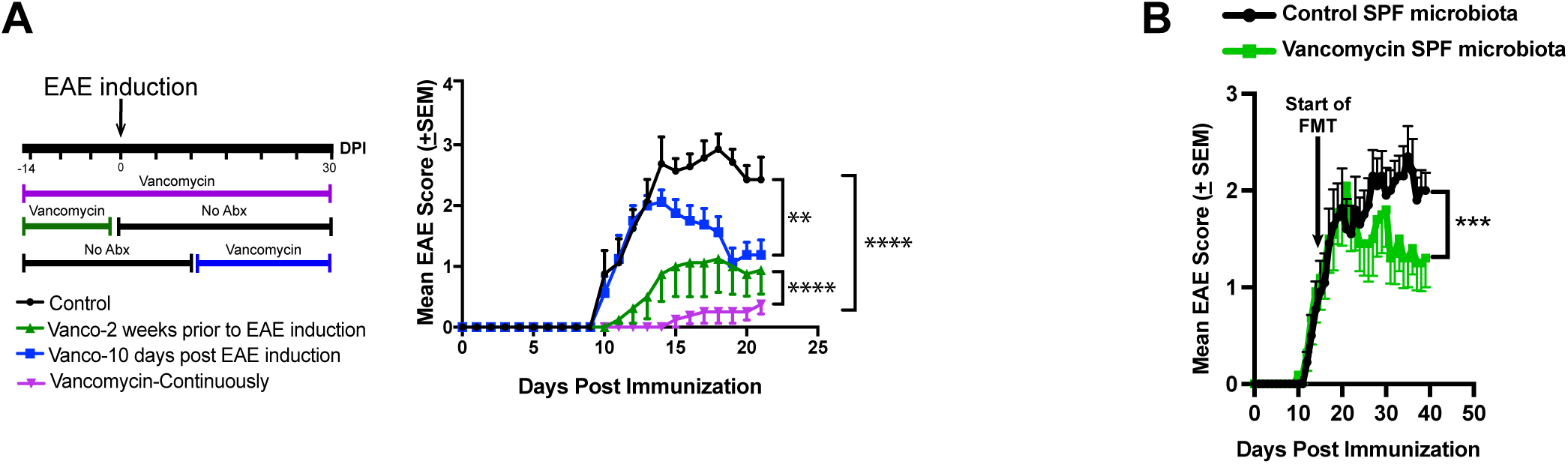
Vancomycin effect on EAE development is mediated via the gut microbiota and is dependent on the length and timing of administration. Conventionally raised mice were given either normal drinking water or vancomycin 0.5mg/mL in drinking water prior to or post immunization with MOG for EAE induction **(A)** Experimental timeline for vancomycin administration (left), and Mean EAE clinical scores over time (right) in untreated and vancomycin-treated mice. Error bars denote mean ± SEM (n = 8-10 mice/group). Representative data of three independent experiments. Data were analyzed using Freidman test with Dunn’s correction for multiple comparisons. *p< 0.05, **p< 0.01, ***p< 0.001, ****p< 0.0001. Conventionally raised mice were immunized with MOG for EAE induction and starting at day 15 post immunization, half of the mice received microbiota from untreated mice and the remaining half received microbiota from vancomycin treated mice daily until the end of the experiment. **(B)** Mean EAE clinical scores over time. Error bars denote mean ± SEM (n = 8- 10 mice/group). Shown are representative data from two independent experiments. Freidman test with Dunn’s correction for multiple comparisons. *p< 0.05, **p< 0.01, ***p< 0.001, ****p< 0.0001. SPF: specific pathogen free.

For experiments characterizing the effects of vancomycin and ampicillin on EAE development (Figure 2A), antibiotics were administered once daily via gavage from day 9 post immunization until the end of the experiment. Mice received 2.5mg of vancomycin (Research Products International Corp.) or 5mg of ampicillin (Research Products International Corp.) daily.

**Figure 2.**
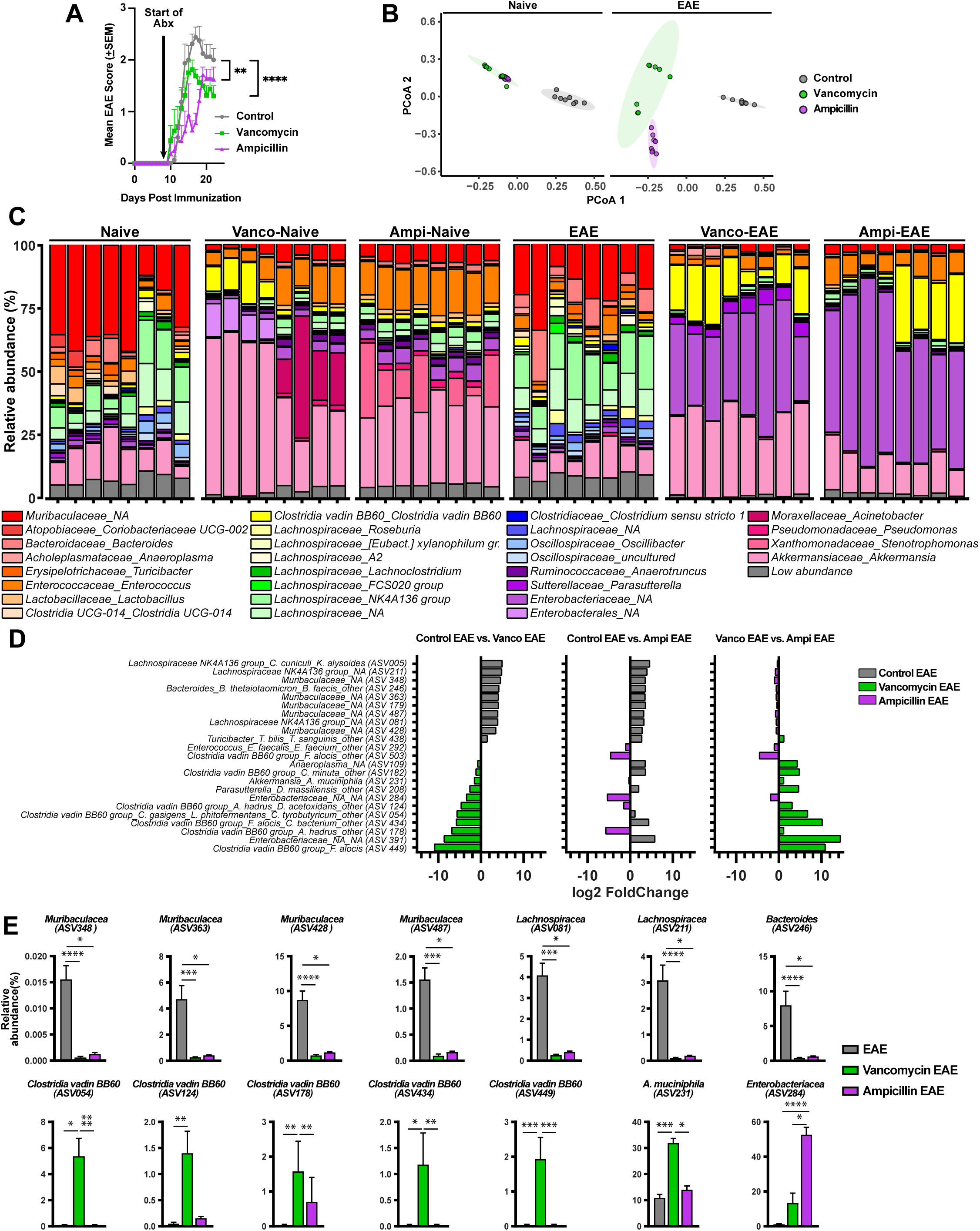
Effect of vancomycin on the gut microbiota during the late stage of EAE. Conventionally raised mice were immunized with MOG for EAE induction and starting at day 9 post immunization, mice were treated with vancomycin, ampicillin or vehicle (water) once daily via oral gavage for the duration of the experiment. **(A)** Mean clinical EAE scores overtime. Error bars denote mean ± SEM (n = 8 mice/group). Representative data of two independent experiments. Freidman test with Dunn’s correction for multiple comparisons. *p< 0.05, **p< 0.01, ***p< 0.001, ****p< 0.0001. **(B)** Principal coordinate analysis of intestinal microbiota samples based on Bray-Curtis showing significantly different clustering of untreated and antibiotic-treated naïve and EAE mice for all pairwise comparisons in naïve and immunized states (q<0.01). Each dot represents a mouse. **(C)** Taxa plots showing compositional differences in fecal microbiota at the genus level between untreated and antibiotic-treated naïve and EAE mice. **(D)** Species level antibiotic-induced changes in the gut microbiome composition of EAE mice. Results are expressed as log2 fold change, two-way ANOVA with false discovery rate. All taxa showing significant differences in relative abundance in any pairwise comparisons are shown. **(E)** Relative abundance of selected bacterial species altered in antibiotic treated and untreated EAE mice. Results are presented as mean ± SEM (n = 8 mice/group). Data were analyzed using the Kruskal-Wallis test, followed by Dunn’s test for multiple comparisons. *p< 0.05, **p< 0.01, ***p< 0.001, ****p< 0.0001. ASV: amplicon sequence variant.

### Fecal microbiota transplantation

Donor mice received either vancomycin 0.5 mg/mL in drinking water or normal drinking water for 2 weeks. Two weeks post vancomycin initiation, ceca were collected under aseptic conditions and processed in an anaerobic chamber to preserve the viability of intestinal bacteria. The content of 1 cecum from vancomycin treated mice, or 3 ceca from control mice were transferred to pre-reduced and anaerobically sterilized PBS (Anaerobe Systems Cat. no. AS-908) and homogenized by vortexing at room temperature. The fecal homogenates were freshly prepared on the day of first gavage. Remaining fecal slurries were aliquoted, stored at −80°C, and thawed only once for subsequent gavages. Recipient C57BL/6J mice were administered via gavage 200 μL of the control- or vancomycin- cecal content homogenate 3 times per week starting at day 15 post EAE induction until the end of the experiment.

### *Lactobacillus reuteri* culture and administration

*Lactobacillus reuteri*, strain CF48-3A (Cat#: HM-102; Biodefense and Emerging Infections Research Resources Repository) was grown in anaerobic conditions at 37 °C in brain-heart infusion (BHI) medium (Cat #: DF0037178; Fisher Scientific). Bacterial cultures were grown for 48-72 hours, centrifuged and resuspended in anaerobic sterile PBS to obtain a suspension of live bacteria at a density of OD600 = 1.3. Mice were administered 200 μL of bacteria suspension via oral gavage 3 times a week beginning either 3 weeks prior to EAE induction or 16 days post MOG immunization, until the end of the experiment. Control mice for this experiment received sterile anaerobic PBS (vehicle) via oral gavage.

### Blood collection

Mice were restrained manually by scruffing the skin of the dorsal neck. A 5mm lancet (Fisher, Cat# NC9891620) was used to puncture the facial vein on the right side of the face. Blood was allowed to drip freely into a microtainer tube blood collection with lithium heparin (Fisher, Cat#13-680-62). Hemostasis was achieved by using clean, dry gauze applied directly to the access site. Plasma was separated from cellular fraction by centrifugation and subsequently stored in a −80°C freezer.

### Tissue collection

Immediately after blood collection, mice were anesthetized with 5% isoflurane and intracardially perfused through the left ventricle with ice-cold HBSS. Spleens, spinal cords, distal colons and ceca were removed and processed immediately or stored in conditions appropriate for downstream applications. Ceca and distal colons were snap frozen and stored at −80°C for metabolomics and 16S rRNA gene sequencing respectively. Spleens were collected for total CD4 T cells isolation and spinal cords were collected in ice cold HBSS for astrocytes isolation.

### 16S rRNA gene sequencing and analysis

Distal colons were collected as described above. Bacterial DNA was isolated from distal colons using MoBio PowerLyzer PowerSoil Kit (Qiagen). DNA concentration was quantified using the Quant-iT dsDNA Assay Kit, broad range (Life Technologies) and normalized to 2 ng/μl on a Beckman handling robot. Amplicon preparation and sequencing was performed as previously described^18^. Briefly, bacterial 16S rDNA PCR including no template controls were set up in a separate PCR workstation using dual-indexed primers. PCR reactions contained 1 µM of each primer, 4 ng DNA, and Phusion Flash High-Fidelity PCR Master Mix (Thermo Fisher Scientific). Reactions were held at 98°C for 30 s, proceeding to 50 cycles at 98°C for 10 s, 45°C for 30 s, and 72°C for 30 s and a final extension of 2 min at 72°C. Amplicons were evaluated by gel electrophoresis. The sequencing library was prepared by combining equivolume amounts of each amplicon, size-selected and concentrated using AMPure XP beads (0.8X, Beckman). Library concentration was quantified by Qubit and qPCR, mixed with 15% PhiX, diluted to 4 pM and subjected to paired-end sequencing (Reagent Kit V2, 2x150bp) on an Illumina MiSeq sequencer. Taxonomy was assigned using the Scikit-learn classifier^19^ pre-trained on the Silva 132 99% OTUs 515-806 region (silva-132-99-515-806-nb-classifier). Taxonomy summary plots were generated based on relative abundance at the genus level. Beta diversity was estimated using Bray-Curtis, and distances were then used to perform principal coordinate analysis as implemented in QIIME. Differences in beta diversity were tested using PERMANOVA with FDR correction for multiple comparison testing. Compositional differences in the microbiome were determined using a two-way ANOVA with false discovery rate.

### Metabolomics

Ceca and plasma samples were shipped to Metabolon, Inc. (Morrisville, NC) for the targeted analysis of bile acids and dietary tryptophan and tyrosine derived metabolites, respectively. Metabolite levels were analyzed using a liquid chromatography-tandem mass spectrometry (LC-MS/MS) system (Agilent 1290 UHPLC/Sciex QTrap 5500).

### Isolation of astrocytes from adult mouse spinal cord

Astrocytes were sorted as previously described^15^ and outlined in Supplementary Fig. 4. Briefly, spinal cords were subjected to homogenization using a tissue homogenizer. Subsequently, cells were pelleted and resuspended in 70% Percoll. An overlay of 37% Percoll was then added on top of 70% Percoll and cells mixture, after which samples were centrifuged for 30 minutes at 800 G at 25 °C. Mononuclear CNS cells were collected from the 70–30% interphase, further centrifuged and resuspended in FACS buffer (HBSS + 2% FBS + 1M HEPES + 0.5M EDTA). Samples were incubated with anti-mouse CD32 for 5 min at 4°C to block Fc receptors and stained with fluorochrome conjugated antibodies to CD45R/B220 PE (1:200) (clone RA3-6B2, BD Bioscience #553089), Ter119 PE (1:200) (Biolegend #116207), Olig4 PE (1:200) (R&D Systems #FAB1326P), CD105 PE (1:200) (clone MJ7/18, Invitrogen #12105182), CD140a PE (1:200) (clone APA5, Invitrogen #12140181), Ly6G PE (1:200) (clone 1A8, Biolegend #127608), CD11b PE-Cy7 (1:200) (clone M1/70, BD Bioscience #561098), Ly6C PerCp (1:200) (clone HK1.4, Biolegend #128028), CD45 APC (1:200) (clone 30-F11, Invitrogen #17045183), CD11c APC-Cy7 (1:200) (clone HL3, BD Bioscience #561241) for 20 minutes at 4°C in the dark. After staining, cells were washed and incubated with 7-AAD viability dye (Biolegend #420403) for 20 minutes at 4°C in the dark to exclude dead cells. Following additional washes, cells were resuspended in FACS buffer and astrocytes were sorted using a BD CSM5L sorter after negative selection of B cells, T/NK cell subset, erythroid cells, oligodendrocytes, endothelial cells, mesenchymal cells, monocytes, granulocytes, neutrophils, microglia, hematolymphoid cells and dendritic cells. We confirmed that sorted cells expressed astrocyte specific markers *Gfap*, *Aldh1l1* and *Aqp4* (**Fig S4G**). Using a separate set of sorted astrocytes and microglia, we confirmed significantly higher expression of GFAP and Aldh1l1 in astrocytes compare to microglia via qPCR.

### Bulk RNA sequencing

#### Antibiotics study

Ultra-low input RNA-seq library preparation, HiSeq sequencing and analysis were conducted at Azenta US, Inc (South Plainfield, NJ, USA). Ultra-low input RNA sequencing library was prepared by using SMART-Seq HT kit for full-length cDNA synthesis and amplification (Takara, San Jose, CA, USA), and Illumina Nextera XT (Illumina, San Diego, CA, USA) library was used for sequencing library preparation. Briefly, cDNA was fragmented, and an adaptor was added using Transposase, followed by limited-cycle PCR to enrich and add index to the cDNA fragments. The sequencing library was validated on the Agilent TapeStation (Agilent Technologies, Palo Alto, CA, USA), and quantified by using Qubit 2.0 Fluorometer (ThermoFisher Scientific, Waltham, MA, USA) as well as by quantitative PCR (KAPA Biosystems, Wilmington, MA, USA). The sequencing libraries were multiplexed and clustered on a flowcell. After clustering, the flowcell was loaded on the Illumina HiSeq instrument according to the manufacturer’s instructions. The samples were sequenced using a 2x150 Paired End (PE) configuration. Image analysis and base calling were conducted by the HiSeq Control Software (HCS). Raw sequence data (.bcl files) generated from Illumina HiSeq was converted into fastq files and de-multiplexed using Illumina’s bcl2fastq 2.17 software. One mismatch was allowed for index sequence identification. After investigating the quality of the raw data, sequence reads were trimmed to remove possible adapter sequences and nucleotides with poor quality using Trimmomatic v.0.36. The trimmed reads were mapped to the Homo sapiens reference genome available on ENSEMBL using the STAR aligner v.2.5.2b. BAM files were generated as a result of this step. Unique gene hit counts were calculated by using feature Counts from the Subread package v.1.5.2. Only unique reads that fell within exon regions were counted.

After extraction of gene hit counts, the gene hit counts table was used for downstream differential expression analysis. Using DESeq2, a comparison of gene expression between the groups of samples was performed. The Wald test was used to generate P values and Log2 fold changes. Genes with adjusted P-values < 0.05 and absolute log2 fold changes >1 were considered differentially expressed genes for each comparison.

#### Lactobacillus study

Ultra-low input RNA-seq library preparation and sequencing reactions were conducted at the Broad Institute (Cambridge, MA, USA) as previously described^20^. Demultiplexed fastq files reflecting RNA-seq data were trimmed for low quality using Trimmomatic^21^ and filtered for artificial adapter sequences, followed by application of STAR^22^ for alignment to the reference mouse transcriptome (vGRCm38) and gene level quantification using RSEM^23^. Normalization of gene abundance profiles across the sample set was performed using DE-Seq2^24^, followed by differential expression analysis of gene families (Wald Test with False Discovery Rate p-value adjustment) to identify differentially expressed genes between groups of interest.

### Quantitative PCR

Total CD4 T cells were isolated from spleens using CD4 beads from Miltenyi (Cat# 130-117-043, Miltenyi). Total RNA was isolated from CD4 T cells using the PicoPure RNA extraction kit (Cat# KIT0204, Thermo Fisher) following manufacturer’s instructions. First strand cDNA synthesis was performed for each RNA sample for up to 1μg of total RNA using a High-Capacity cDNA Reverse Transcription kit (Fisher, Cat# 4374966). Quantitative PCR (qPCR) analysis was performed in an Applied Biosystems ViiA 7 system (Applied Biosystems), using the SYBR Green master mix and specific primers for IL17A (FWD: agcaagagatcctggtcctgaa, RV: catcttctcgaccctgaaagtga), IL10 (FWD:agaagcatggccctgaaatcaagg RV: cttgtagacaccttggtcttggag), and IFN-g (FWD: gccatcagcaacaacataagcgtc, RV: ccactcggatgagctcattgaatg). GAPDH (FWD: ACAGTCCATGCCATCACTGCC, RV: GCCTGCTTCACCACCTTCTTG) was used as endogenous control to normalize for differences in the amount of total RNA across samples. The relative gene expression was calculated using the comparative CT method^25^.

### Quantification and statistical analysis

Spearman’s rank correlation coefficient was calculated across pairwise elements in different partitions using the spearman r function from the stat module in ScipPy. p-values were corrected for experiments of interest using Python’s False-Discovery Rate Benjamini-Hochberg (FDR-BH) function from the statsmodels package. All graphs and additional statistical analyses were performed using the GraphPad Prism software (GraphPad Software, San Diego, CA, USA). Mann-Whitney test was used for comparisons between two groups. Kruskal-Wallis followed by Dunn’s test was used for comparisons between 3 or more groups. EAE clinical scores were analyzed over time with the nonparametric Friedman test with Dunn’s correction for multiple comparisons. Details about the statistical tests, sample sizes, and p-values are indicated in each figure.

## RESULTS

### Vancomycin effect on EAE development depends on time of administration

One study showed that administration of vancomycin during the late stage of EAE had no effect on disease severity or recovery^15^. We have previously reported that administration of vancomycin for 2 weeks prior to EAE induction ameliorates EAE^13^. To examine if this apparent discrepancy of vancomycin effect on EAE development from these two studies is due to difference in timing of administration, we next conducted experiments to investigate how the timing and length of administration of vancomycin affect EAE progression. C57BL/6J (B6) EAE mice were administered vancomycin at various time points spanning pre- and post- EAE induction (**Fig 1A**). We observed that mice receiving vancomycin continuously starting at 2 weeks prior to EAE induction (purple group) did not develop disease (**Fig 1A**). Mice who were treated with vancomycin daily for 2 weeks (green group) prior to EAE induction developed mild disease consistent with our prior study^13^ (**Fig 1A**). Mice who received vancomycin daily starting at the time of symptom onset (day 10 post immunization; blue group) developed disease severity that was comparable to that seen in untreated mice (black group) but had better disease recovery that was similar to that seen in mice treated with vancomycin prior to EAE induction (**Fig 1A**). Hence, vancomycin can both prevent and treat disease in EAE mice. We have shown that germ-free mice colonized with microbiota from vancomycin treated mice developed less severe disease than mice colonized with microbiota from conventionally raised mice^13^. To determine if the microbiota derived from vancomycin treated mice can attenuate established EAE, EAE mice either received microbiota from untreated mice or vancomycin treated mice starting at day 15 post immunization (after symptoms onset). We found that mice fed microbiota from vancomycin treated mice had better disease recovery than mice fed microbiota from conventionally raised mice (**Fig 1B**). Hence, the neuroprotective effects of vancomycin are mediated via the microbiota. Taken together, our results revealed that vancomycin neuroprotective effects depend on the length and timing of administration.

### Taxa belonging to Clostridia vadin BB60 group *are enriched in* mice treated with vancomycin during the late stage of EAE

As mentioned above, gut commensals derived from vancomycin treated mice ameliorate disease recovery in EAE mice. Hence, to identify bacteria modulating disease recovery during EAE, we conducted 16S rRNA gene sequencing to investigate the gut microbiota composition during the late stage of EAE in untreated, vancomycin and ampicillin treated EAE mice. EAE was induced in B6 mice via MOG immunization and at day 9 post immunization (pre-symptoms onset; early stage EAE) these EAE mice were divided in the following 3 groups: 1 group was administered vancomycin once daily, the second group received ampicillin once daily and the third group received vehicle once daily via oral gavage until the end of the experiment. As expected, we found that vancomycin treated mice had better recovery than the untreated EAE mice (**Fig 2A**). Ampicillin treated EAE mice had delayed symptom onset compared to untreated and vancomycin treated mice and displayed better disease recovery than untreated EAE mice (**Fig 2A**). At day 23 (late stage EAE), mice were sacrificed and stool samples were used for 16S rRNA gene sequencing. PCoA plots revealed that the untreated, vancomycin and ampicillin treated EAE mice have a distinct microbiota composition (q<0.01 for all pairwise comparisons; **Fig 2B**). We observed increased abundance of *Muribaculaceae* family, *LachnospiraceaeNK4A136 group*, *Bacteroides* and *Turicibacter* genera in untreated EAE mice compared to vancomycin and ampicillin treated EAE mice (**Fig 2C-E**). We found that six taxa belonging to *Clostridia vadin BB60 group*, *Enterobacteriaceae family*, *Akkermansia*, *Parasuterella* and *Anaeroplasma* genera were enriched in vancomycin treated mice compared to untreated and ampicillin treated EAE mice (**Fig 2C-E**). Interestingly, the family *Muribaculaceae* has been shown to positively correlate with serum levels of the pro-inflammatory cytokine IL-1β^26^. Furthermore, we have previously shown that *Akkermansia* ameliorates EAE^12,27^. Interestingly, a study reported that the *Clostridia vadin BB60 group* is depleted in treatment naïve MS patients compared to treated MS patients and healthy controls^28^. Hence, our findings suggest that vancomycin administration post EAE induction leads to the depletion of pro-inflammatory bacteria and proliferation of neuroprotective bacteria with a predominance of taxa belonging to *Clostridia vadin BB60 group* genus.

### Vancomycin alters metabolism of bile acids, tryptophan and tyrosine derived metabolites during the late stage of EAE

As mentioned above, we found that vancomycin treatment of EAE mice induced significant change in the gut microbiota composition leading to improved disease recovery. A prior study reported that vancomycin-resistant bacteria are indole-producers and that 3 indole derivatives namely, indole-3-propionic acid (3-IPA), indoxyl-3-sulfate (3-IS) and indole-3-aldehyde, are important modulators of the inflammatory response during the recovery phase of EAE^15^. Hence, we next performed serum metabolomics to assess if vancomycin induced changes in the gut microbiota composition are linked to changes in serum levels of indole derivatives. We found decreased serum levels of 3-IS, 3-IPA in vancomycin and ampicillin treated naïve and EAE mice compare to untreated naïve and EAE mice (**Fig 3A-B**). We observed increased serum levels of 3-IPA in untreated EAE mice compared to untreated naïve mice (**Fig 3B**). Vancomycin treated EAE mice had increased serum 3-IS level compared to vancomycin treated naïve mice as well as ampicillin treated mice (**Fig 3B-C**). 3-ILA levels were comparable in untreated, vancomycin and ampicillin treated naïve mice (**Fig 3A**). We observed increased serum levels of 3-ILA in vancomycin treated EAE mice compared to vancomycin treated naïve mice as well as untreated and ampicillin treated EAE mice (**Fig 3A-C**). Serum tryptophan levels were not affected by antibiotic treatment or EAE induction (**Fig 3A-C**). Hence, our results showed that vancomycin depleted bacteria producing 3-IS, 3-IPA but promoted the proliferation of bacteria producing 3-ILA. Interestingly, a prior study reported that high salt diet exacerbates EAE at least in part by inducing the depletion of 3-ILA producing *Lactobacillus* species^29^. In addition, another study reported that ILA ameliorates EAE^30^. Consistent with our findings, one study observed increased levels of 3-IS in the cerebrospinal fluid of untreated MS patients compared to treated MS patients and they found that 3-IS is neurotoxic^31^. A prior study found that IPA ameliorates disease recovery in ampicillin treated EAE mice^15^. We observed that vancomycin treated EAE mice had significantly lower serum levels of IPA compared to untreated EAE mice, suggesting that IPA is not required to attenuate neuroinflammation during EAE.

**Figure 3.**
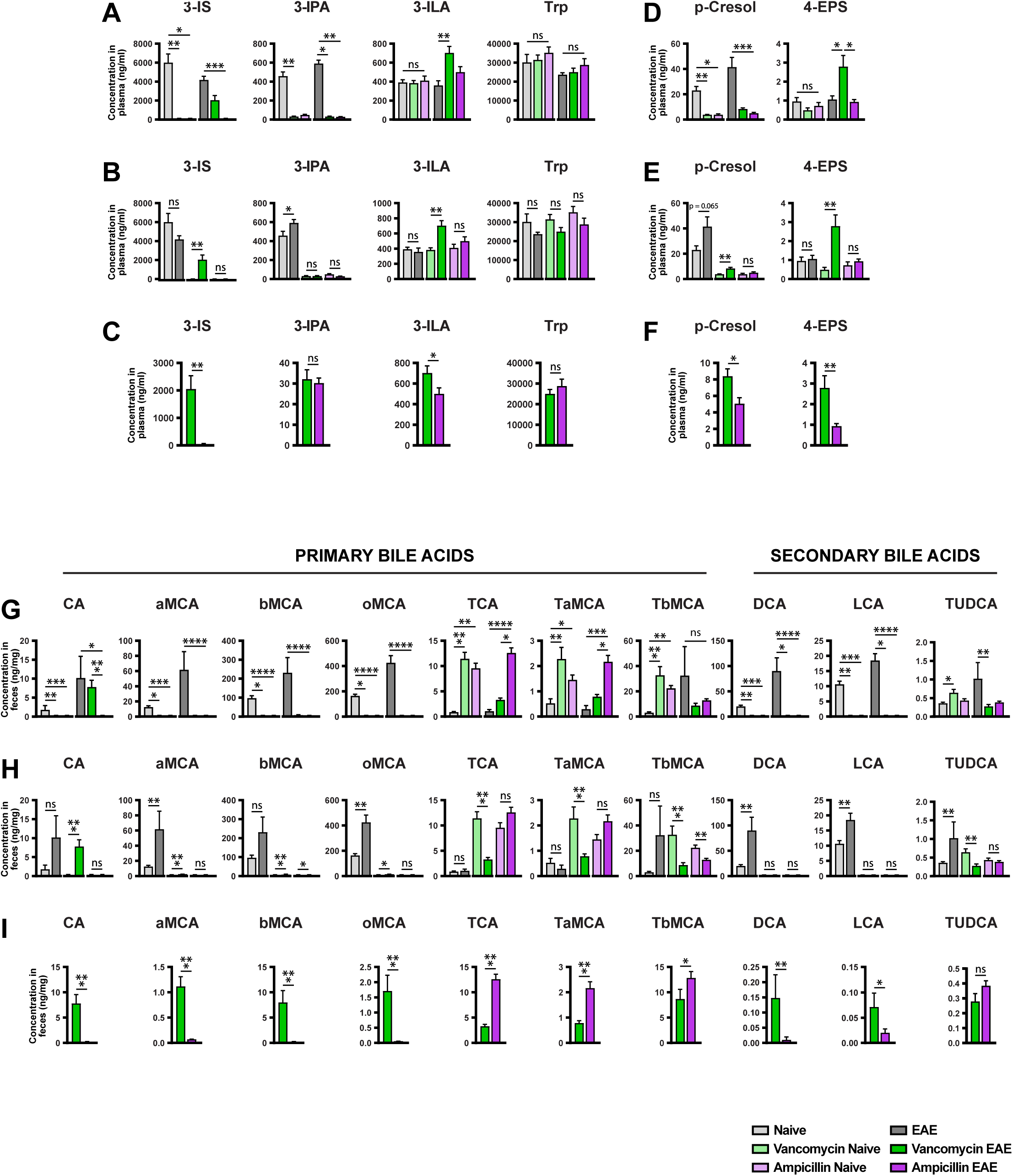
Effect of vancomycin on serum and stool levels of metabolites during the late stage of EAE. Plasma and ceca samples collected from untreated and antibiotic-treated naïve and EAE mice at day 23 post immunization were used for metabolites detection and quantification by liquid chromatography-tandem mass spectrometry. **(A-C)** levels of tryptophan-derived metabolites and **(D-F)** levels of tyrosine-derived metabolites. **(G-I)** levels of bile acids. Results are presented as mean ± SEM (n = 6-8 mice/group). Data were analyzed using the Kruskal-Wallis test with the Dunn’s correction (A, B, D, E, G, H, J), or the Mann-Whittney non-parametric test (C, F, I). *p< 0.05, **p< 0.01, ***p< 0.001, ****p< 0.0001.

In addition to tryptophan derived metabolites, tyrosine derived metabolites, p-cresol, and 4-EPS have been implicated in MS/EAE^31,32^ and impaired neuronal myelination^33,34^. Hence, we also examined the effect of vancomycin on serum levels of p-cresol and 4-EPS in our mice. We found decreased serum levels of p-cresol in vancomycin and ampicillin treated naïve and EAE mice compared to untreated naïve and EAE mice (**Fig 3D**). We observed increased serum levels of p-cresol in untreated EAE mice compared to untreated naïve mice (**Fig 3E**). 4-EPS serum level was comparable in untreated, vancomycin and ampicillin treated naïve mice (**Fig 3D**). 4-EPS serum levels were increased in vancomycin treated EAE mice compared to vancomycin treated naïve mice as well as untreated and ampicillin treated EAE mice (**Fig 3D-F**). Serum 4-EPS levels were low in all mice groups and to date, no study has examined the effect of 4-EPS in EAE mice or MS patients. Prior studies have shown that p-cresol is pro-inflammatory and neurotoxic^31,35^. Consistent with these findings, we observed that vancomycin depleted gut commensals that produce p-cresol. Taken together, our findings suggest that a putative mechanism by which vancomycin ameliorates disease recovery in EAE mice is via inducing the proliferation of anti-inflammatory 3-ILA producing bacteria and depleting bacteria producing the neurotoxins 3-IS and p-cresol.

Another study reported altered levels of primary and secondary bile acids in MS patients^36^. That same study also reported that supplementation of EAE mice with the secondary bile acid metabolite TUDCA ameliorated disease by modulating glial cells function during the recovery phase. Hence, we next examined the effect of vancomycin on stool levels of bile acids. We found that mice treated with vancomycin or ampicillin had very low stool levels of most primary bile acids compared to untreated mice (**Fig 3G-I, Fig S1A-C**). Vancomycin and ampicillin treated naïve mice had increased stool levels of primary bile acids TaMCA, TbMCA and TCA compared to untreated naïve mice (**Fig 3G-H**). Stool levels of TaMCA, TbMCA and TCA were decreased in vancomycin treated EAE mice compared to vancomycin treated naïve mice and ampicillin treated EAE mice (**Fig 3H-I**). Stools levels of TaMCA, TbMCA and TCA were comparable in untreated, and vancomycin treated EAE mice (**Fig 3G**). We found that mice treated with vancomycin or ampicillin had very low stool levels of all secondary bile acids (**Fig 3G-I**). We observed a slight increase in stool levels of TUDCA in vancomycin treated naïve mice compared to untreated naïve mice (**Fig 3G**). However, stool TUDCA levels were higher in untreated EAE mice compared to untreated naïve mice as well as vancomycin and ampicillin treated EAE mice (**Fig 3G-H**). Furthermore, vancomycin treatment increased stool levels of TUDCA in naïve mice but had no effect on TUDCA levels in EAE mice (**Fig 3G-H**). Taken together, our results suggest that elevated stool TUDCA could play a role in disease recovery in untreated EAE mice but not in vancomycin treated EAE mice. We found that untreated EAE mice had increased stool levels of DCA compared to untreated naïve mice (**Fig 3G-H**). Vancomycin and ampicillin treated mice had negligible stool levels of DCA (**Fig 3G-I**). Interestingly, prior studies have reported that DCA is pro-inflammatory in a mouse model of colitis^37–39^. Hence, our results suggest that depletion of DCA producing bacteria could be another mechanism accounting for vancomycin neuroprotective effect during the recovery phase of EAE.

### Correlation between metabolites and bacteria abundance during the late stage of EAE

Indole derivatives, p-cresol, and 4-EPS are gut commensals derived products. We found that vancomycin modulates the gut microbiota composition as well as serum levels of indole derivatives, p-cresol and 4-EPS in EAE mice. To assess if vancomycin induced changes in the gut microbiota composition of EAE mice are linked to serum levels of tryptophan and tyrosine metabolites, we performed Spearman correlation between serum levels of these metabolites and bacterial abundance. We observed that 3-IS and P-cresol positively correlated with two taxa within the *Lachnospiraceae NK4A136* group (ASV 81, ASV211), *Bacteroides* (ASV246) and four *Muribaculaceae* taxa (ASV348, ASV363, ASV428 and ASV 487) (**Fig 4A-C; Fig S2**). Interestingly, these seven taxa were decreased in vancomycin treated EAE mice compared to untreated EAE mice (**Fig 2D-E**). We found that 3-IS and p-cresol were negatively correlated with a taxa within the *Enterobacteriaceae* family (ASV284) (**Fig 4A-C**). In addition, *Enterobacteriaceae* taxa (ASV284) was enriched in vancomycin treated EAE (**Fig 2D-E**). ILA and 4-EPS positively correlated with 4 *Clostridia vadinBB60 group* genera (ASV 124, ASV178, ASV434 and ASV449) and *Akkermansia municiphila* (ASV 231) (**Fig4A, 4D-E; Fig S2**). These five taxa were increased in vancomycin treated EAE mice compared to untreated EAE mice (**Fig 2D-E**). We observed that ILA negatively correlated with four *Muribaculaceae* taxa (ASV348, ASV363, ASV428 and ASV 487) (**Fig 4A, 4D; Fig S2)** that were depleted in vancomycin treated EAE mice (**Fig 2D-E**) and positively correlated with 3-IS and p-cresol (**Fig 4B-C; Fig S2**). Consistent with the finding that 3-IS and p-cresol are neurotoxic^31^, we observed that vancomycin treatment depleted bacteria linked with increased serum levels of p-cresol and 3-IS and promoted the proliferation of bacteria linked with decreased serum levels of these metabolites. Consistent with prior studies reporting that ILA has anti-inflammatory properties and ameliorates EAE^29,30^, we observed that vancomycin treatment promoted the proliferation of bacteria linked with increased serum levels of 3-ILA.

**Figure 4.**
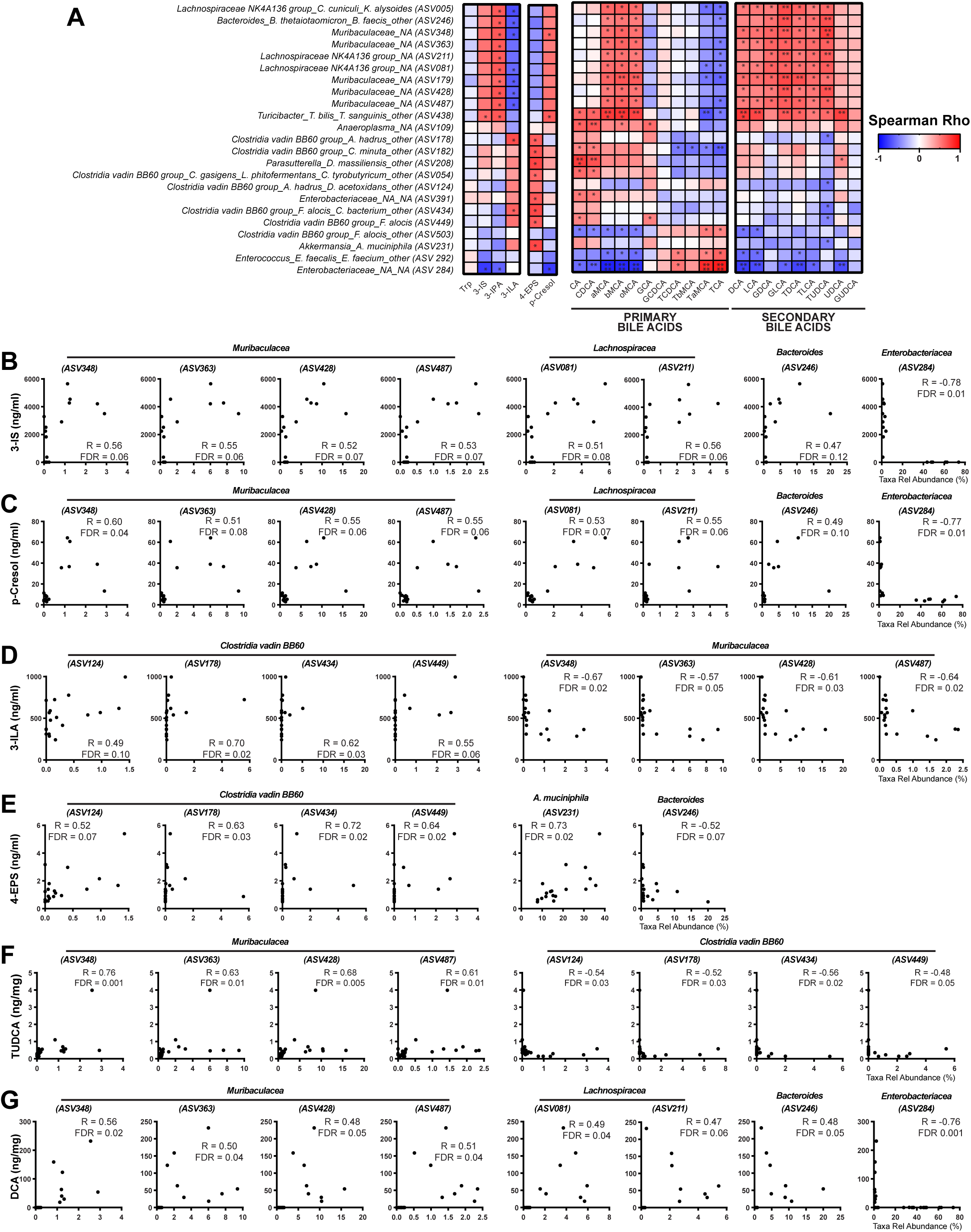
Microbiota associated with metabolites during the late stage of EAE. Spearman’s correlation between relative abundance of indicated taxa and metabolite concentrations. **(A)** Correlation matrix of selected taxa at the lowest classifiable levels showing significant correlations (FDR<0.05). *p< 0.05; **p< 0.01; ***p< 0.001; ****p< 0.0001. **(B-G)** Scatter plots of selected taxa at the lowest classifiable levels showing significant correlations. ASV: amplicon sequence variant, R: Spearman’s correlation coefficient.

As mentioned above, vancomycin treated mice had very low stool levels of bile acids compared to untreated mice. Given that secondary bile acids originate from the gut microbiota, we next ran a Spearman correlation to examine if vancomycin induced changes in the gut microbiota composition of EAE mice are linked to stool levels of bile acids. We observed that stool levels of TUDCA and DCA (two secondary bile acids that have been implicated in autoimmune diseases) positively correlate with five taxa belonging to the *Muribaculaceae* family (ASV179, ASV348, ASV363, ASV428 and ASV487), three taxa belonging to the *Lachnospiraceae NK4A136 group* (ASV005, ASV81, ASV211) and one taxa belonging to *Bacteroides* (ASV246) (**Fig 4A, 4F-G; Fig S2**). Interestingly, these taxa also positively correlate with serum p-cresol and/or 3-IS levels (**Fig 4B-C; Fig S2**) and were enriched in untreated EAE mice but depleted in vancomycin treated EAE mice (**Fig 2D-E**). Consistent with prior studies showing that DCA is a pro-inflammatory bile acid^37–39^, we found that vancomycin treatment depleted bacteria linked with increased stool levels of DCA.

### Astrocytes from vancomycin treated mice exhibit distinct transcriptomes during late stage of EAE

Since CD4^+^ T cells are key players in the pathogenesis of the EAE model we tested the effects of vancomycin and ampicillin on CD4^+^ T cell cytokine production and noted no direct anti-inflammatory effects of vancomycin and ampicillin on CD4^+^ T cells (**Fig S3**). These findings suggest that vancomycin and ampicillin do not appear to mediate their effects on EAE through alteration of T cell function.

Prior studies have reported that astrocytes derived pathways are important modulators of neuroinflammation during the recovery phase of EAE^15^. Furthermore, indole derivatives and the secondary bile acid TUDCA have been shown to modulate astrocyte function during the recovery phase of EAE^15,36^. We observed that vancomycin treatment initiated in the early stage of EAE (prior to symptoms onset) improved disease recovery and altered serum levels of indole derivatives and stool levels of secondary bile acids. Hence, to assess how vancomycin induced changes in levels of indoles and secondary bile acids affect astrocyte function, we analyzed mRNA expression in astrocytes isolated from the spinal cord of these EAE mice by RNA sequencing (RNA Seq). We detected 17,690 expressed genes of which 3,422 were differentially expressed among all mice groups (**Fig S4-5A**).

We found that vancomycin treatment altered the expression of 1414 genes in astrocytes from EAE mice compared to untreated EAE mice. 1030 genes were differentially expressed in astrocytes from vancomycin treated EAE mice versus ampicillin treated EAE mice. Astrocytes isolated from vancomycin treated EAE mice showed decreased levels of pro-inflammatory genes including Tspo (**Fig 5A, 5G**) as well as increased expression of genes with antimicrobial activity like Cxcl14^40^ (**Fig 5A, 5G**). We also observed increased expression of the pro-inflammatory chemokine, Ccl12 in vancomycin treated EAE mice compared to untreated or ampicillin treated EAE mice (**Fig 5G**). Previous studies have suggested a potentially protective role for the pro-inflammatory chemokine, Ccl2, in multiple sclerosis, with elevated levels observed during remission compared to relapses^41^. This increase in Ccl2 expression during remission suggests that anti-inflammatory monocytes may be recruited to inflamed areas, thus contributing to repair and remyelination processes^42^. Therefore, it is reasonable to postulate that increased level of Ccl12 expression in vancomycin treated EAE mice may similarly facilitate the recruitment of anti-inflammatory monocytes, particularly during the recovery phase. A study showed that indole derivatives: 3-IS, 3-IPA and indole-3-aldehyde binding to Ahr on astrocytes induces anti-inflammatory pathways suppressing neuroinflammation during the recovery phase of EAE^15^. We observed increased Ahr expression in astrocytes from ampicillin treated EAE mice (**Fig 5B-C, 5G**). Furthermore, ampicillin treated EAE mice had high 3-ILA serum levels whereas serum levels of 3-IS and 3-IPA were negligible (**Fig 3A-C**). Vancomycin treatment did not change Ahr expression on astrocytes (**Fig 5A-C, 5G**) but we observed increased serum levels of indole-3-lactic acid in vancomycin treated EAE mice compared to untreated and ampicillin treated EAE mice (**Fig 3B-C**). Taken together, our results suggest that the neuroprotective effects of vancomycin and ampicillin treatments may be in part mediated via activation of Ahr on astrocytes by 3-ILA.

**Figure 5.**
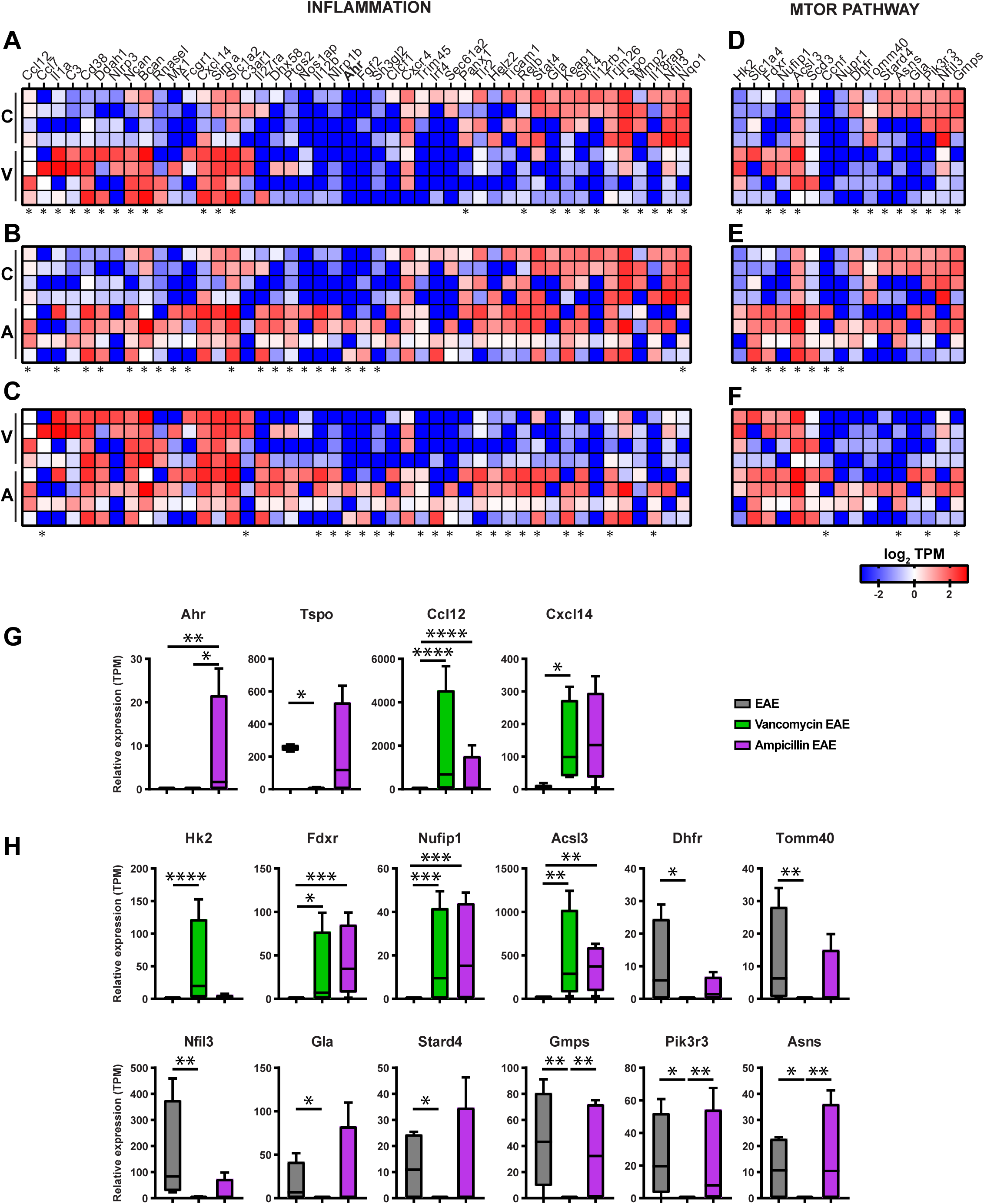
Vancomycin treatment regulates the expression of genes associated with inflammation and the mTOR pathway in astrocytes during the late stage of EAE. Total RNA was extracted from astrocytes isolated from spinal cords of naïve and EAE mice treated with Vancomycin, Ampicillin or vehicle. Differentially expressed (**A-C**) inflammatory genes and (**D-F**) mTOR pathway genes across treatment groups were identified using DESeq2 and analyzed using the Wald test. Genes with adjusted p-values < 0.05 and absolute log2 fold changes >1 were defined as differentially expressed genes in each pairwise comparison. Individual values are shown (n = 4 mice/group). Results are expressed as log 10 TPM (Transcripts per kilobase million). *p< 0.05, Wald test. **G.** Selected differentially expressed inflammatory genes; *p< 0.05; **p< 0.01; ***p< 0.001. **H.** Differentially expressed mTOR pathway genes; Results are presented as median with interquartile ranges, whiskers represent minimum and maximum values *p< 0.05; **p< 0.01; ***p< 0.001, Wald test.

Another group of genes in astrocytes whose expression was modulated by vancomycin belongs to the mTOR pathway. We found that vancomycin treatment induced increased expression of Hk2, Fdxr, Nufip1 and Acsl3 in astrocytes of EAE mice compared to untreated EAE mice (**Fig 5D-F, 5H**). We observed that vancomycin treatment induced downregulation of Dhfr, Tomm40, Stard4, Asns, Gla, Pik3r3, Nfil3 and Gmps compared to untreated EAE mice (**Fig 5D-F, 5H**).

### Correlation between mTOR genes, bacteria abundance and metabolites during late stage of EAE

Prior studies have shown that indole derivatives and the secondary bile acid TUDCA modulate astrocyte function during the recovery phase of EAE. Furthermore, we found that vancomycin treatment modulates gut microbiota composition, serum levels of indole derivatives, p-cresol, 4-EPS and stool levels of secondary bile acids. We also observed that vancomycin induced changes in the expression of many genes in astrocytes including those belonging to the mTOR pathway. To determine if there is a link between level of expression of mTOR pathway genes in astrocytes, gut commensals abundance and serum levels of indole derivatives, p-cresol, 4-EPS and secondary bile acids, we performed Spearman correlation. Fdxr positively correlated with *Enterobacteriaceae* taxa (ASV284) and negatively correlated with 3-IS, P-cresol, DCA, LCA and TDCA (**Fig. 6A-B**). Nufip1 negatively correlated with *Turicibacter* genus (ASV438), 3-IS, p-cresol and TDCA. Ascl3 negatively correlated with 2 *Lachnospiraceae NK4A136 group* taxa (ASV005, ASV211), p-cresol, DCA, LCA, GLCA, TDCA, TLCA (**Fig. 6A-B)**. Fdxr, Nufip and Ascl3 were upregulated in vancomycin treated EAE mice compared to untreated EAE mice (**Fig 5A, 5H**). Genes upregulated in astrocytes from vancomycin treated EAE mice are linked with decreased serum levels of 3-IS, p-cresol, DCA and TDCA (**Fig 6B**).

**Figure 6.**
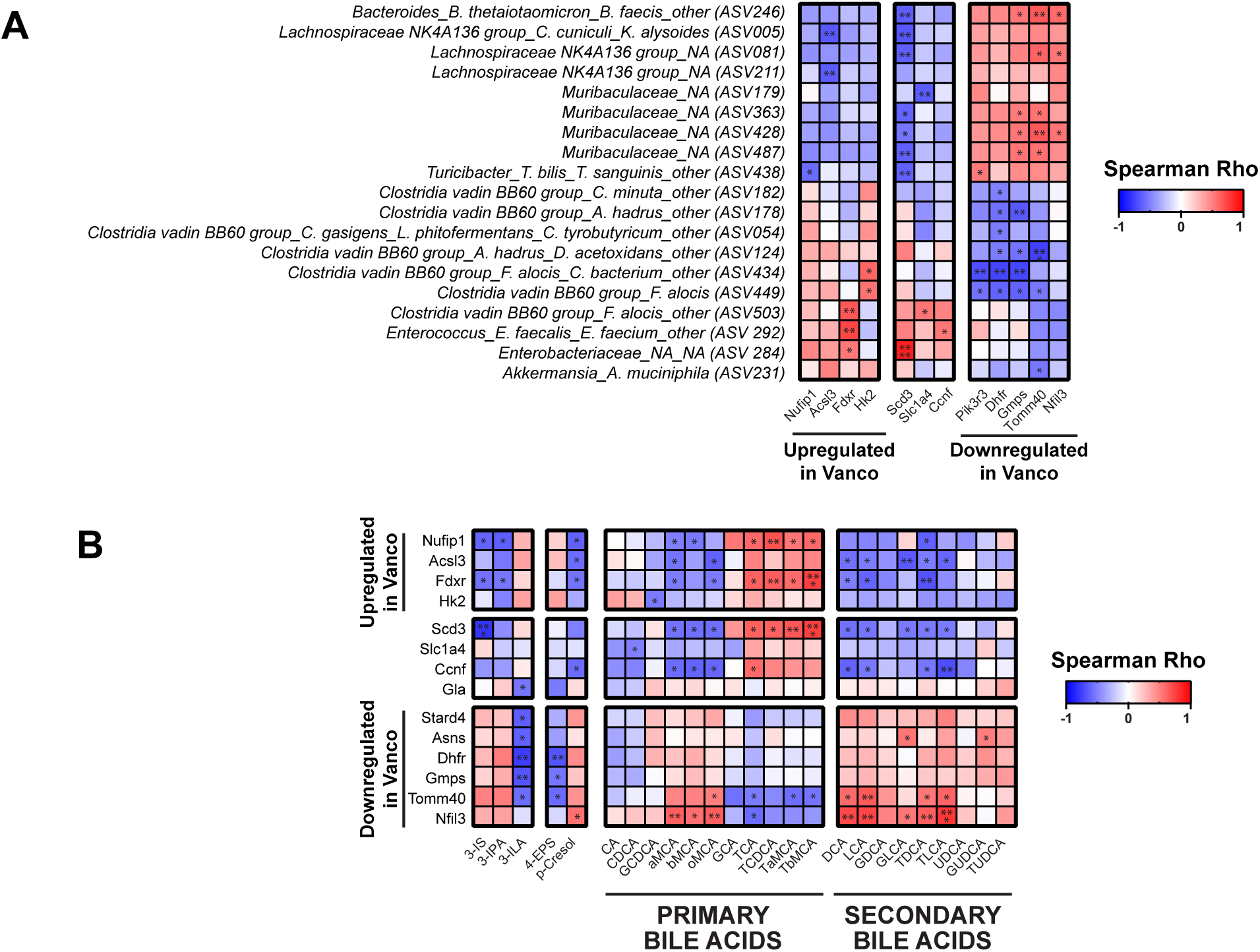
Microbiota and metabolites linked with level of expression of mTOR pathway genes in astrocytes during the late stage of EAE. **(A)** Spearman’s correlation between differentially expressed mTOR pathway genes and abundance of selected taxa. **(B)** Spearman’s correlation between differentially expressed mTOR pathway genes and metabolite concentrations. *p< 0.05; **p< 0.01; ***p< 0.001; ****p< 0.0001. Taxa names are indicated at the lowest classifiable level. ASV: amplicon sequence variant.

Dhfr negatively correlated with 6 taxa belonging to *Clostridia vadin BB60 group* (ASV54, ASV124, ASV178, ASV182, ASV434, ASV449), 3-ILA and 4-EPS (**Fig. 6A-B**). Tomm40 positively correlated with *Lachnospiraceae NK4A136 group* (ASV81), ASV246, 3 taxa belonging to *Muribaculaceae* family (ASV363, ASV428, ASV487), DCA, LCA, TDCA, TLCA and negatively correlated with 2 taxa belonging to *Clostridia vadin BB60 group* (ASV124, ASV449), *Akkermansia municiphila* (ASV231), 3-ILA and 4-EPS (**Fig. 6A-B**).

Nfil3 positively correlated with *Lachnospiraceae NK4A136 group* (ASV81), *Bacteroides* genus (ASV246), *Muribaculaceae* (ASV428), DCA, LCA, GLCA, TDCA, TLCA and p-cresol (**Fig. 6A-B**). Gmps positively correlated with *Bacteroides* genus (ASV246), 3 *Muribaculaceae* taxa (ASV363, ASV 428, ASV 487) and negatively correlated with 4 *Clostridia vadin BB60 group* taxa (ASV124, ASV178, ASV434, ASV449), 3-ILA and 4-EPS (**Fig. 6A-B**). Dhfr, Tomm40, Nfil3 and Gmps were downregulated in vancomycin treated EAE mice compared to untreated EAE mice (**Fig 5D-F, 5H**). Genes downregulated in astrocytes from vancomycin treated EAE mice are linked with either increased levels of 3-ILA and 4-EPS or decreased levels of p-cresol, DCA and TDCA.

### Lactobacillus reuteri modulate mTOR pathway genes expression in astrocytes during late stage of EAE

Our correlation studies suggest that vancomycin modulates the expression of genes belonging to the mTOR pathway at least in part by promoting the proliferation of ILA-producing bacteria. Taxa belonging to the genus *Clostridia vadin BB60 group* showed a positive correlation with serum ILA levels in EAE mice. To date, species belonging to genus *Clostridia vadin BB60 group are not commercially available and* have been poorly characterized hence, it is not known if they are indole producers.

Several *Lactobacillus* species are known indole producers and given that we have previously observed increased abundance of *Lactobacillus* species in vancomycin treated mice^14^, we examine the relative abundance of *Lactobacilli* in our mice. We observed increased relative abundance of 4 *Lactobacillus* species in vancomycin and ampicillin treated naïve mice (**Fig 7A**). We have previously observed increased abundance of *L. reuteri*, an ILA-producing gut commensal, in vancomycin treated EAE mice^14^. Consistent with these findings, we found that *L. reuteri* was increased in vancomycin and ampicillin treated naïve mice compared to untreated naïve mice (**Fig 7A**). Of note, *L. reuteri* was below the threshold of detection in EAE mice but this does not mean that it was not present (**Fig 7A).** Hence, it is conceivable that *L. reuteri* was one of the ILA-producing bacteria in EAE mice treated with vancomycin and ampicillin. Prior studies have reported that *L. reuteri* ameliorates EAE^43,44^ and three other studies reported that *L. reuteri* exacerbates EAE^35,45,46^. Hence, to determine how *L. reuteri* affects EAE development in our environment, B6 mice were gavaged with *L. reuteri* pre-EAE induction (green group) or at day 16 post immunization (black group). We found that administration of *L. reuteri* ameliorates disease recovery (**Fig 7B**).

**Figure 7.**
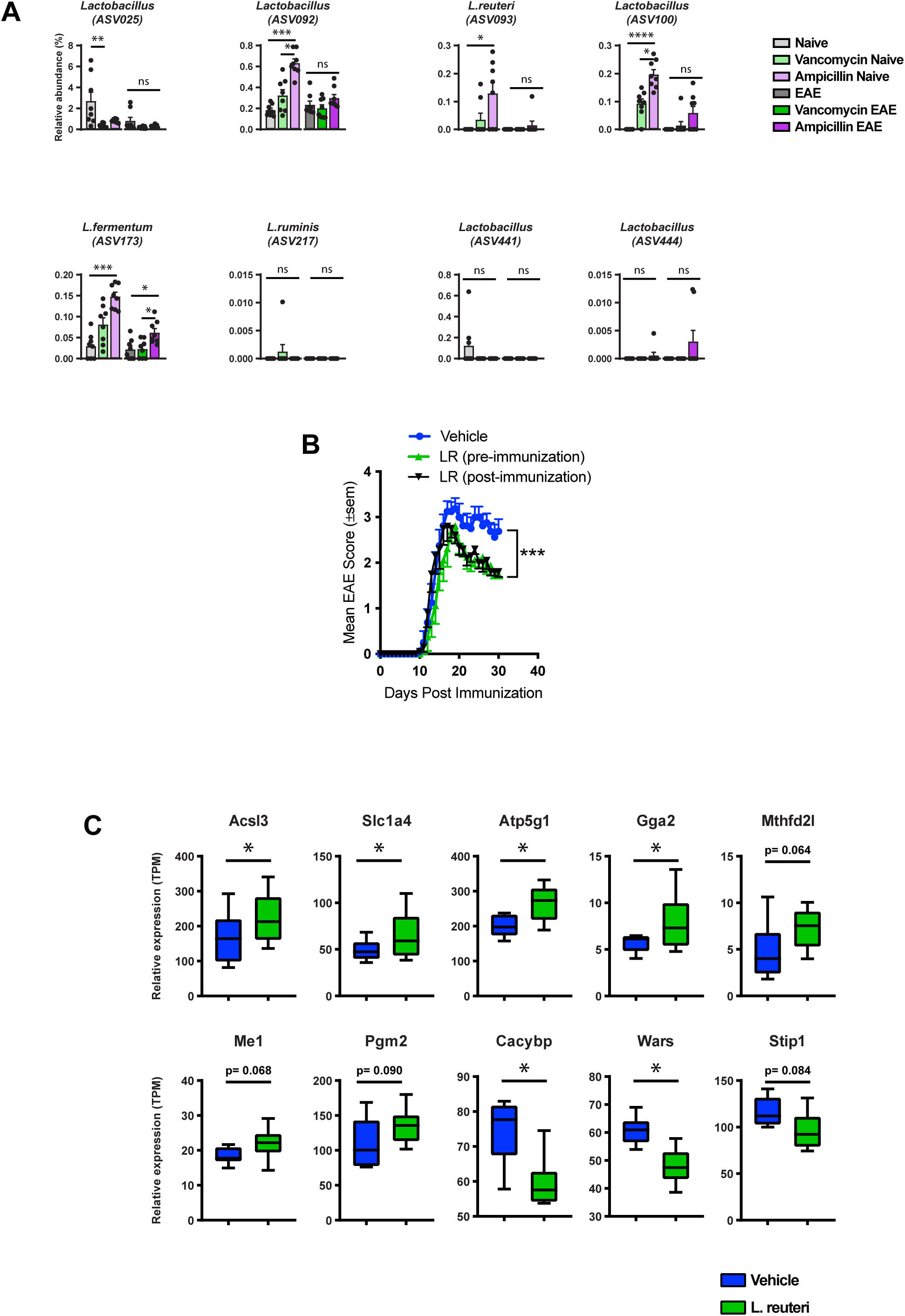
*Lactobacillus reuteri* ameliorates EAE and modulates the expression of the mTOR pathway genes in astrocytes. **(A)** Relative abundance of eight ASVs belonging to the genus *Lactobacillus* in untreated and antibiotic-treated naïve and EAE mice. Error bars denote mean ± SEM with individual values (n = 8 mice/group). Data were analyzed using the Kruskal-Wallis test, followed by Dunn’s test for multiple comparisons. Taxa names are indicated at the lowest classifiable level. **(B)** Mice were orally gavaged 3 times a week with vehicle (blue), or *Lactobacillus reuteri* starting either 3 weeks before (green), or 15 days after (black) EAE induction, until the end of the experiment. **(B)** Mean EAE clinical scores over time. Error bars denote mean ±SEM (n=10 mice/group). Representative data of three independent experiments. Data were analyzed using the Freidman test with Dunn’s correction for multiple comparisons. **(C)** mTOR pathway genes that are differentially expressed between mice treated with *L. reuteri* and vehicle were identified using DESeq2 and analyzed using the Wald test. Genes with p-values < 0.05 were defined as differentially expressed genes. Results are expressed as TPM (Transcripts per kilobase million). Boxplots show median with interquartile ranges, whiskers represent minimum and maximum values (n = 4-6 mice/group). *p< 0.05, **p< 0.01, ***p< 0.001, ****p< 0.0001, ASV: amplicon sequence variant.

Given that we observed a correlation between ILA levels and the expression of several genes belonging to the mTOR pathway, we next performed RNA sequencing to examine the effect of *L. reuteri* on mTOR gene pathway expression in astrocytes. RNA sequencing revealed 22,128 expressed genes of which 1,118 were differentially expressed among all mice groups (**P<0.05; Fig S4-5B**). We found that *L. reuteri* regulated the expression of 10 genes belonging to the mTOR pathway in astrocytes (**Fig 7C**). We observed increased expression of Acsl3 and Slc1a4 in astrocytes from *L. reuteri* treated mice compared to mice who received vehicle (**Fig 7C**). Interestingly, Acsl3 expression was increased in vancomycin and ampicillin treated EAE mice compared to untreated EAE mice (**Fig 5D-E**). Slc1a4 expression was increased in ampicillin treated mice (**Fig 5E**). Ascl3 expression in vancomycin treated mice negatively correlated with stool levels of DCA. Interestingly, we observed decreased stool DCA levels in mice fed *L. reuteri* compared to mice receiving vehicle (data not shown). Taken together, these results suggest that increased ILA:DCA ratio in vancomycin treated mice and *L. reuteri* fed mice leads to increased Acsl3 expression.

## DISCUSSION

Over the past several years, many pre-clinical and clinical studies have gathered findings supporting a role of the gut microbiota in the pathophysiology of MS. Notably, a prior study showed that binding of indole derivatives, which are gut commensals derived tryptophan metabolites, on the AhR receptor in astrocytes triggered pathways that suppress neuroinflammation during the late stage of EAE^15^. These findings suggest that indole-producing bacteria modulate astrocyte function however, the identity of these bacteria remains obscure. Here we show that oral administration of vancomycin to EAE mice during the early stage of EAE (prior to symptoms onset) ameliorate disease recovery. We observed an increase in the abundance of gut commensals belonging to the *Clostridia vadin BB60 group* in vancomycin treated EAE mice compared to untreated and ampicillin treated EAE mice. Interestingly, one study reported decreased abundance of *Clostridia vadin BB60 group* in treatment naïve MS patients compared to treated MS patients or healthy subjects^28^. Another study reported depletion of *Clostridia vadin BB60 group* in EAE mice compared to naïve mice starting at 8 days post immunization^44^. These findings suggest that *Clostridia vadin BB60 group* may have anti-inflammatory properties. Consistent with this notion, two studies have reported that *Clostridia vadin BB60 group* is protective in IBD^47,48^.

Given prior reports that indoles derivatives and the secondary bile acid, TUDCA, modulate astrocyte function during the chronic phase of EAE, we next examined the effect of vancomycin on serum levels of these metabolites. We found increased serum levels of ILA and decreased IPA, 3-IS and p-cresol in vancomycin treated EAE mice compared to untreated EAE mice. Interestingly, ILA has been shown to have anti-inflammatory properties^29,30^. Furthermore, another study reported that ILA is decreased in progressive MS patients compared to relapsing-remitting MS (RRMS) patients^32^. Consistent with these findings, another group identified ILA as one of the most depleted serum metabolites in MS compared to controls^49^. Interestingly, we found a positive correlation between serum levels of ILA and abundance of several taxa belonging to *Clostridia vadin BB60 group*. Hence, these data suggest that *Clostridia vadin BB60 group* may produce ILA and this could account for its anti-inflammatory effect. Future studies in gnotobiotic mice are warranted to characterize the interaction between taxa belonging to *Clostridia vadin BB60 group* and the host during EAE. One study reported that 3-IPA and 3-IS ameliorate disease during the chronic phase of EAE^15^. However, another study showed that several tryptophan metabolites are pro-inflammatory and exacerbate EAE^35^. Furthermore, one study showed that 3-IS and p-cresol are neurotoxins^31^. Consistent with these findings, another study reported increased levels of p-cresol in progressive MS patients compared to RRMS^32^. We observed decreased stool levels of DCA in vancomycin treated EAE mice compared to untreated EAE mice. To date, no study has examined the effect of DCA in EAE mice. Previous studies have reported that DCA exacerbates inflammation in a mouse model of colitis^37–39^. Hence, our results suggest that vancomycin treatment improved disease recovery by eliminating neurotoxins (3-IS and p-cresol) producing or pro-inflammatory metabolites (DCA) producing gut commensals and triggering proliferation of bacteria that produce anti-inflammatory metabolites (ILA). Of note, we found that stool bile acids levels were increased in untreated EAE mice compared to untreated naïve mice. Notably, we observed increased stool TUDCA in untreated EAE mice compared to naïve untreated mice. Given prior report that TUDCA ameliorates disease recovery in EAE mice^36^, elevated TUDCA level in untreated EAE mice could be a host response to suppress neuroinflammation thus allowing mice to enter the recovery phase of the disease. Of note, even though vancomycin increased stool level of TUDCA in naïve mice, it had no effect on TUDCA levels in EAE mice.

Given prior reports that indole derivatives and TUDCA modulate astrocyte function^15,36^, we next examine the effect of vancomycin on astrocytes transcriptional profile. We found that vancomycin treatment modulates the expression of several genes belonging to the mTOR pathway. Interestingly, prior studies have shown that inhibiting the mTOR pathway ameliorates EAE via inducing T regulatory cells or suppressing Th1 and/or Th17 cells^50,51^. However, to date, no study has examined the role of the mTOR pathway in astrocytes during EAE. Nonetheless, emerging data indicate that mTOR signaling in astrocytes contributes to key processes, such as endolysosomal remodeling^52^, which could impact neuroinflammation and disease progression. We observed upregulation of Fdxr, Acsl3 and Nufip1 genes belonging to the mTOR pathway in vancomycin treated EAE mice compared to untreated EAE mice. Interestingly, one study reported that a naturally-occuring mouse model with homozygous Fdxr mutations displayed optic atrophy as well as atrophy of the cerebellum and cerebral cortex^53^. They also observed increase in markers of both neurodegeneration and glial activation in these Fdxr mutant mice^53^. These findings suggest that loss of function of Fdxr leads to neuronal loss. Hence, the increased in Fdxr expression that we observed in our vancomycin treated EAE mice compared to untreated EAE mice could have neuroprotective effects. Future studies using astrocyte specific Fdxr KO mice are needed to elucidate the function of this gene during EAE. The functions of Acsl3 and Nufip1 in astrocytes remain unknown. Interestingly, we found that mTOR pathway genes that are upregulated in astrocytes from vancomycin treated mice had a positive correlation with a taxa enriched in vancomycin treated EAE mice, *Enterobacteriaceae* taxa (ASV284), and a negative correlation with taxa enriched in untreated EAE mice namely *Muribaculaceae* and *Lachnospiraceae NK4A136 group*. We also observed that mTOR pathway genes that are upregulated in astrocytes from vancomycin treated mice negatively correlate with levels of p-cresol, 3-IS and DCA. mTOR pathway genes that were downregulated in astrocytes from vancomycin treated EAE mice had a negative correlation with taxa enriched in vancomycin treated mice, *vadin BB60 group*, and/or a positive correlation with taxa enriched in untreated EAE mice, *Muribaculaceae* and *Lachnospiraceae NK4A136 group*.

mTOR genes that are downregulated in astrocytes from vancomycin treated mice negatively correlate with serum levels of ILA (metabolite enriched in vancomycin treated mice) but positively correlate with levels of 3-IS, p-cresol and DCA (metabolites enriched in untreated EAE mice). Interestingly, *Lactobacilli reuteri*, an ILA producing bacteria, was enriched in vancomycin treated mice. We found that EAE mice fed *L. reuteri* had improved disease recovery compared to mice receiving vehicle. Consistent with this finding, prior studies have reported that *L. reuteri* ameliorates EAE^43,44^. In addition, mice that received *L. reuteri* had altered gene expression of mTOR pathway genes in astrocytes notably a gene upregulated in vancomycin treated EAE mice, Ascl3, was also upregulated in *L. reuteri* fed mice. Ascl3 expression was associated with decreased DCA stool level in *L. reuteri* fed mice as well as vancomycin treated EAE mice. Hence, it could be that high ILA levels lead to decreased DCA levels which then leads to increased Ascl3 expression. Consistent with this notion, one study reported that high levels of the anti-inflammatory metabolite, butyrate, is associated with low levels of DCA^54^. Hence, these results suggest that indole derivatives and DCA can modulate the expression of mTOR pathway genes in astrocytes (**Fig 8**: hypothesized model diagram). Other studies have reported that indole derivatives as well as secondary bile acids regulate the mTOR pathway^55–58^. Furthermore, another study reported that *L. reuteri* modulates mTOR expression^59^. The mTOR pathway has been implicated in several inflammatory and neurodegenerative diseases including MS^50,60–63^. Hence, more studies are needed to elucidate the mechanisms by which indoles derivatives and DCA modulate the expression of mTOR pathway genes in astrocytes during EAE.

**Figure 8.**
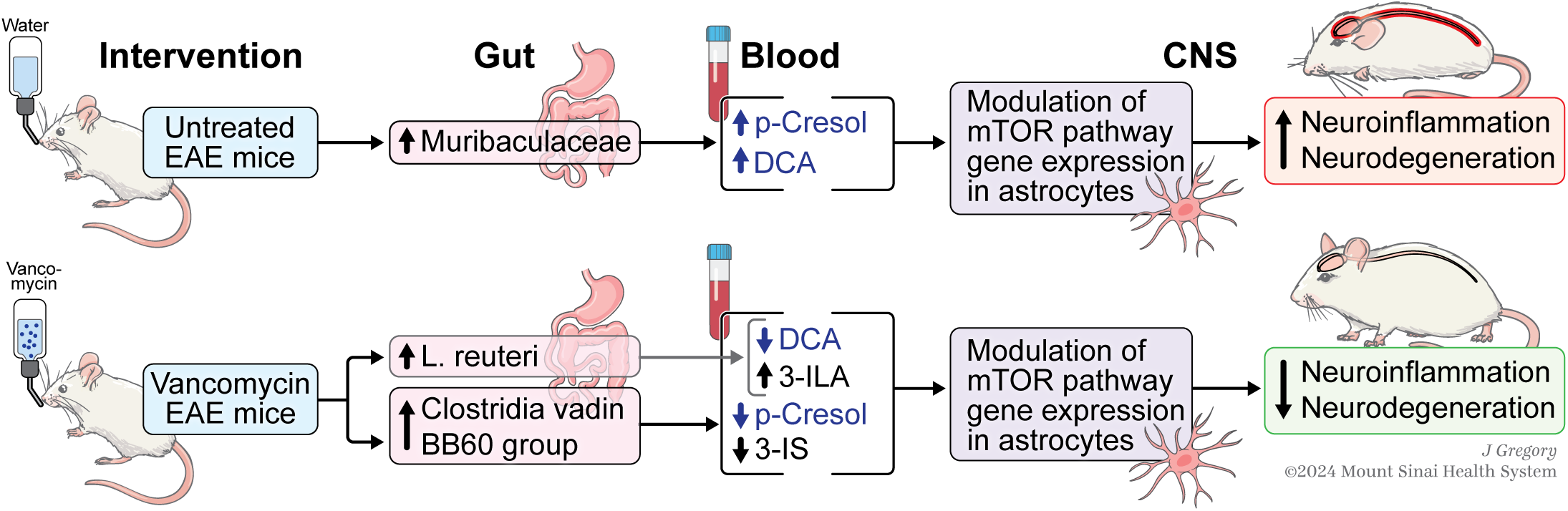
Hypothesized model diagram for this study. Study summary scheme.

Our results also revealed that some indole derivatives as well as secondary bile acids are pro-inflammatory/neurotoxic while others are anti-inflammatory/neuroprotective. Furthermore, even though we and other groups have found 3-IS to be neurotoxic in EAE mice and MS patients^31^, another study reported that this indole derivative is neuroprotective in EAE mice^15^. In addition, contrary to our findings, some studies have observed that strains of *L. reuteri* different from the one used in our study exacerbate EAE^35,45,46^. The conflicting results observed in EAE mice colonized with different strains of *L. reuteri* underscore the complexity of host-microbiome interactions and highlight the need for a personalized medicine approach to probiotic administration. Variations in microbial communities, individual genetic background and environmental factors such as diet that regulate tryptophan metabolism can influence the outcome of *L. reuteri* supplementation by changing the ratio of neurotoxic to neuroprotective metabolites for example. Therefore, integrating metagenomics, metabolomics, host genetics data and environmental factors into clinical assessments could help clinicians identify patients most likely to benefit from supplementation with specific strains of ILA-producing *L. reuteri*.

Our study has some limitations. Although we observed increased abundance of the *Clostridia vadin BB60 group* in vancomycin treated EAE mice and a correlation between this taxa and expression of mTOR pathway genes in astrocytes, we cannot establish a causal effect. Additional studies will be required to determine whether monocolonization with this taxa is sufficient to modulate expression of genes in the mTOR pathway. Furthermore, we observed that vancomycin altered the expression of several genes in the mTOR pathway in astrocytes but their functions during neuroinflammation remain unknown. Our work adds to the growing literature on the importance of the mTOR pathway role in CNS inflammatory and neurodegenerative diseases. Previous studies have shown that rapamycin, a specific inhibitor of mTOR, ameliorates EAE via suppression of T effector cells or inducing T regulatory cells^50^ To date, there are no published studies examining the role of the mTOR pathway in astrocytes during EAE. Hence future studies using astrocytes specific KO of mTOR pathway genes of interest are warranted to elucidate the mechanism by which vancomycin induced changes in astrocytes transcriptional profile regulate neuroinflammation during EAE. Finally, our findings suggest that indole derivatives as well as bile acids modulate mTOR pathway genes expression in astrocytes during EAE. Future studies using AhR and G-protein coupled bile acid receptor astrocytes specific KO should be conducted to investigate the effect of indole derivatives and bile acids on the expression of mTOR pathway genes in EAE mice. In addition, other metabolites not measured in this study could also modulate astrocyte function during EAE.

Despite these limitations, our study showed that vancomycin modulates astrocyte function during the chronic phase of EAE. We also observed that vancomycin induced the production of indole-3-lactic acid, an anti-inflammatory metabolite, that is decreased in progressive MS compared to relapsing-remitting MS patients^32^. Furthermore, another group reports that in addition to inhibiting neuroinflammation in MOG induced EAE mice, ILA also promotes remyelination in the cuprizone EAE mouse model (Larissa Jank et al; ACTRIMS 2024 abstract)^64^. These findings suggest that vancomycin could be beneficial for progressive MS patients. While we are currently running a double-blind, placebo-controlled randomized trial to investigate the clinical impact of vancomycin in newly diagnosed, treatment naive MS patients (NCT05539729), future trials will be required to examine the effect of vancomycin in progressive MS patients with low levels of ILA.

## ABBREVIATIONS

3-ILA: 3 Indole Lactic Acid
3-IPA: 3-Indole Propionic Acid
3-IS: 3-Indoxyl Sulphate
4-EPS: 4-Ethyl Phenyl Sulphate
AhR: Aryl Hydrocarbon receptor
aMCA: Alpha Muricholic Acid
bMCA: Beta Muricholic Acid
CA: Cholic Acid
CDCA: Chenodeoxycholic Acid
CNS: Central Nervous System
DCA: Deoxycholic Acid
EAE: Experimental Autoimmune Encephalomyelitis
GCA: Glycocholic Acid
GCDCA: Glycochenodeoxycholic Acid
GDCA: Glycodeoxycholic Acid
GLCA: Glycolithocholic Acid
GUDCA: Glycoursodeoxycholic Acid
IBD: Inflammatory Bowel Disease
LCA: Lithocholic Acid
MS: Multiple Sclerosis
mTOR: mammalian Target Of Rapamycin
oMCA: Omega Muricholic Acid
p-Cresol: p-Cresol Sulphate
RNA Seq: RNA Sequencing
TaMCA: Tauro Alpha Muricholic Acid
TbMCA: Tauro Beta Muricholic Acid
TCA: Taurocholic Acid
TCDCA: Taurochenodeoxycholic Acid
TDCA: Taurodeoxycholic Acid
TLCA: Taurolithocholic Acid
Trp: Tryptophan
TUDCA: Tauroursodeoxycholic Acid
UDCA: Ursodeoxycholic Acid

## AUTHORS’ CONTRIBUTION

SKT, PB, PL, CGV and EMS performed experiments; EW, PB, GB and JCC analyzed the microbiome data; PB analyzed the RNA-Seq data; SKT, PB and PL generated the figures; and SKT and PB wrote the manuscript with input from all authors. SKT conceived the study, designed the analysis, and finalized the manuscript. The authors read and approved the final manuscript.

## DATA AVAILABILITY

The microbiota 16S rRNA sequence data as well as RNA sequencing data will be submitted to the Mendeley Data Repository and made available to the public once the paper is published.

## ACKNOWLEDGMENTS

We acknowledge the support of the Dean’s Flow Cytometry CORE at the Icahn School of Medicine at Mount Sinai. We would like to thank Dr. Francisco Quintana for providing us with a detailed protocol for isolation of astrocytes from mouse spinal cords.

---

**Supplementary Figure 1.**
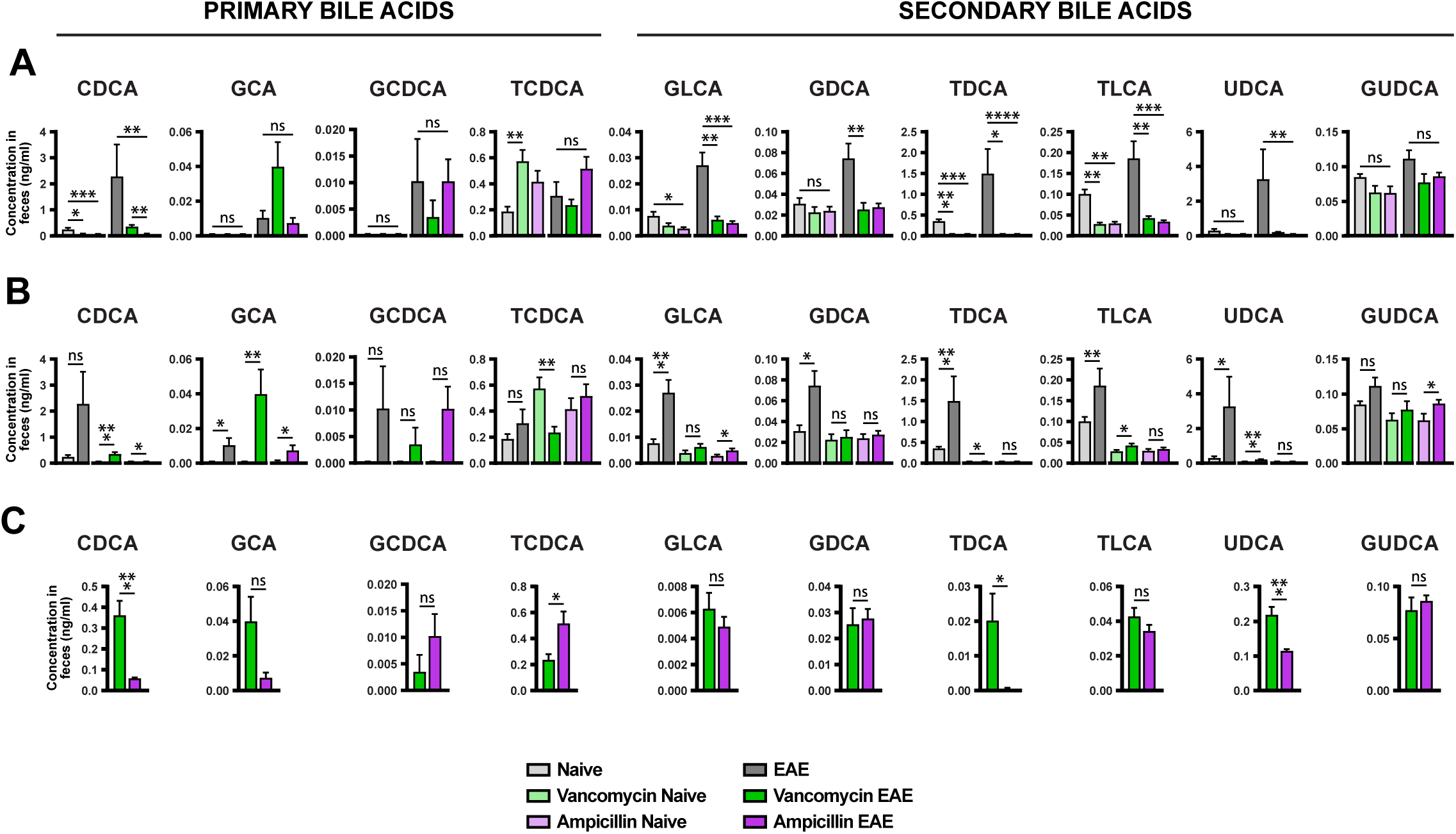
Additional bile acid levels in feces from untreated or antibiotic-treated naïve and EAE mice. (A-C) The concentration of bile acids was measured by liquid chromatography-tandem mass spectrometry in ceca samples. Error bars denote mean ±SEM (n = 8 mice/group). Data were analyzed using the Kruskal-Wallis test with the Dunn’s correction (**A, B**), or the Mann-Whittney non-parametric test (**C**). *p< 0.05, **p< 0.01, ***p< 0.001, ****p< 0.0001

**Supplementary Figure 2.**
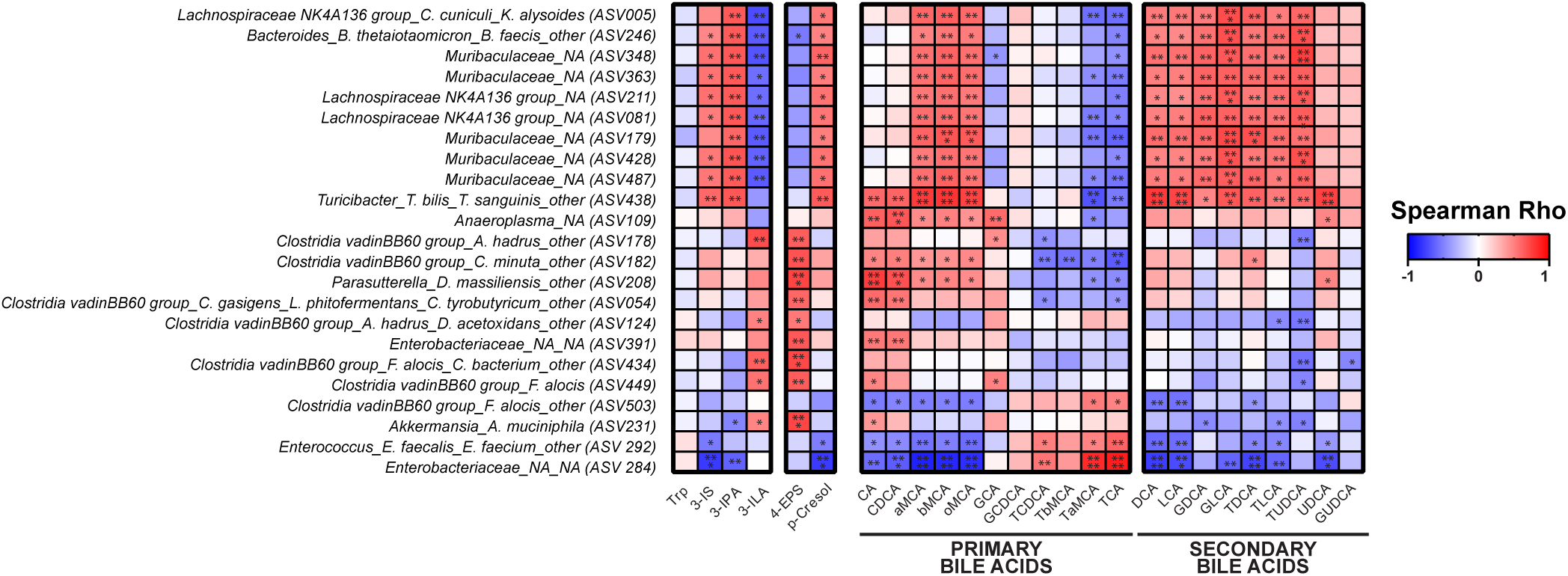
Microbiota associated with metabolites during the late stage of EAE. Spearman’s correlation between relative abundance of indicated taxa and metabolites concentrations. Correlation matrix of selected taxa at the lowest classifiable levels showing significant correlations (non-adjusted p-values). *p< 0.05; **p< 0.01; ***p< 0.001; ****p< 0.0001

**Supplementary Figure 3.**
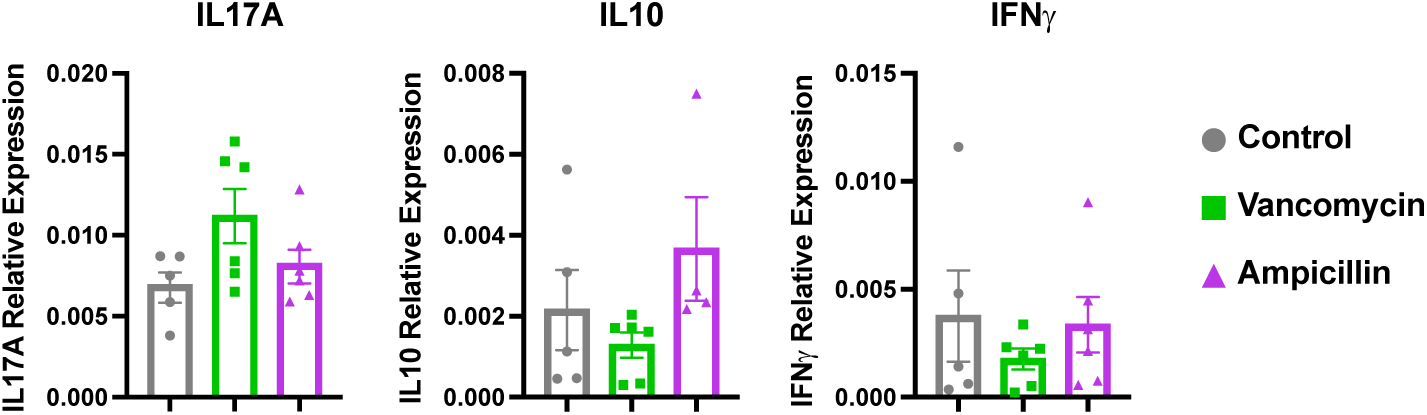
Cytokines expression in spleen. Total CD4^+^ T cells were isolated from spleens using CD4 beads. Relative expression of IL-10, IL-17A and IFN-γ were assessed by qPCR in CD4^+^ T cells from untreated EAE mice as well as EAE mice treated with vancomycin or ampicillin. Representative data of two independent experiments. Results were analyzed using one-way ANOVA followed by Bonferroni’s multiple comparisons test. Error bars denote mean ±SEM with individual values, ns: not significant.

**Supplementary Figure 4.**
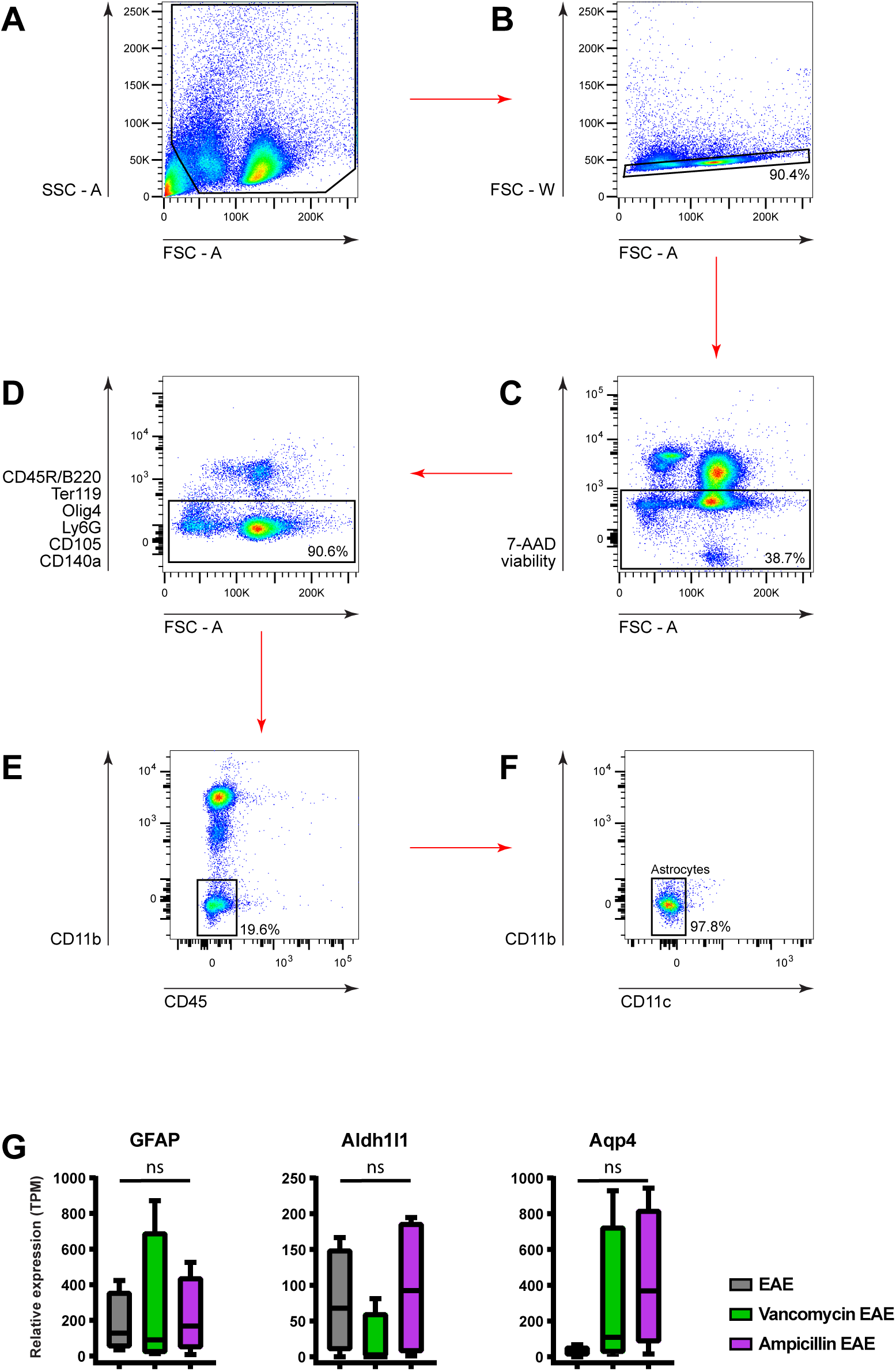
FACS gating strategy for astrocytes sorting. **(A)** Mononuclear CNS cells were gated based on SSC-A versus FSC-A and **(B)** singlets were selected from the FSC-A versus FSC-W dot plot. **(C)** Dead cells were excluded with 7-AAD viability dye. **(D)** Cells positive for CD45R/B220 (B cells, T/NK cell subset), Ter119 (erythroid cells), Olig4 (oligodendrocytes), CD105 (endothelial cells), CD140a (mesenchymal cells), Ly6C (monocytes), Ly6G (granulocytes, neutrophils) were excluded. **(E)** Cells positive for CD45 (hematolymphoid cells) and CD11b (myeloid cells) were excluded. **(F)** Astrocytes were identified as double negative for CD11b and CD11c (dendritic cells). **(G)** We confirmed that we had isolated a relatively pure population of astrocytes by examining the expression of astrocyte-specific markers *Gfap*, *Aldh1l1*, and *Aqp4*. Results are expressed as TPM (Transcripts per kilobase million). Boxplots show median with interquartile ranges, whiskers represent minimum and maximum values (n = 4 mice/group). ns: not significant.

**Supplementary Figure 5.**
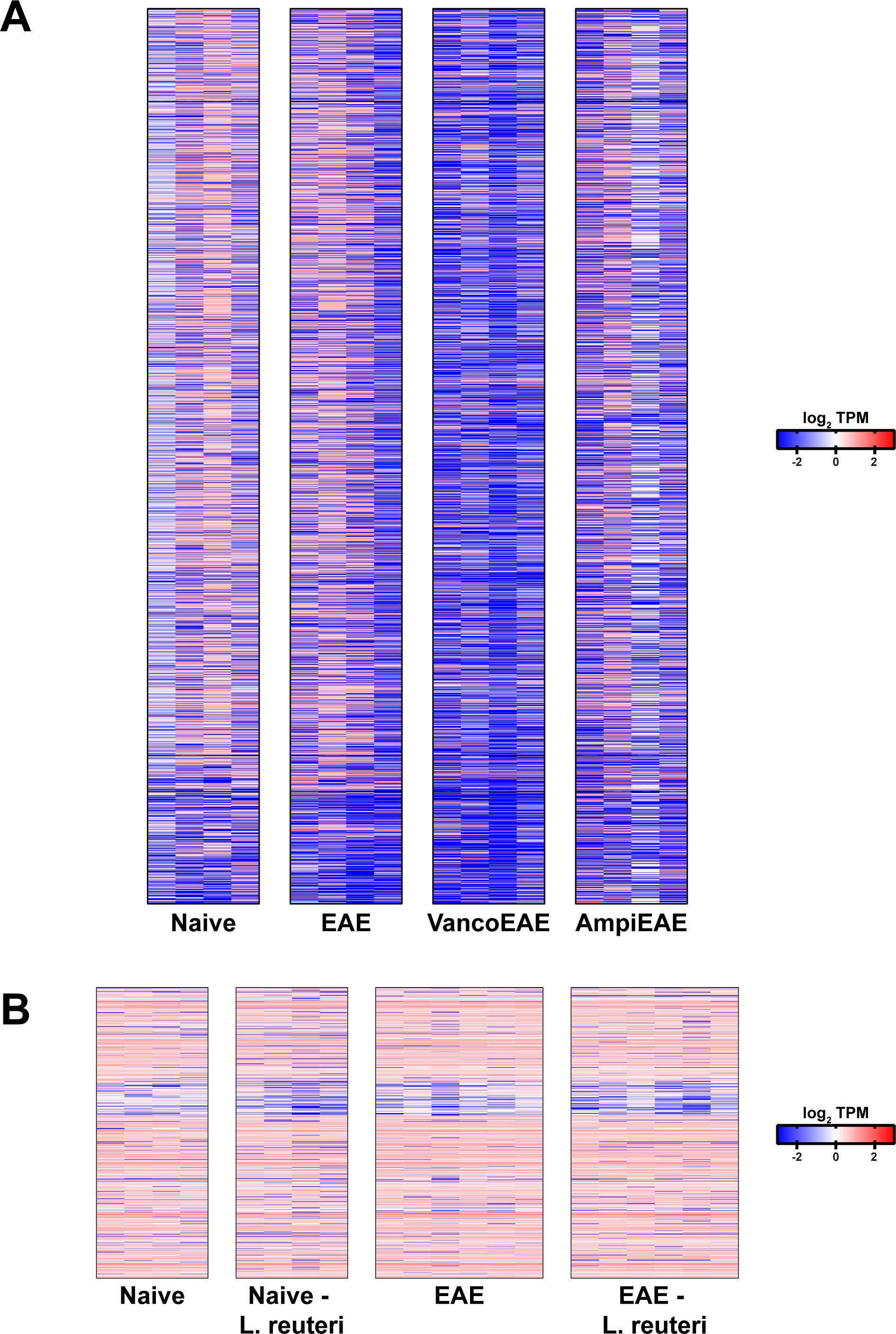
Effect of Vancomycin and *Lactobacillus reuteri* on astrocytes transcriptional profile during the late stage of EAE. **(A)** Heatmaps of 3,422 differentially expressed genes in naïve and untreated or antibiotic treated EAE mice. Pairwise comparisons of gene expression between the treatment groups was performed using DESeq2. Expression data was analyzed using the Wald test. Genes with adjusted p-values < 0.05 and absolute log2 fold changes >1 were defined as differentially expressed genes for each pairwise comparison. All genes differentially expressed in any pairwise comparison are included. The color scale represents the log10 (average TPM + 0.01) value. Individual values are shown (n = 4mice/group) **(B)** Heatmaps of 1,118 differentially expressed genes between untreated mice and mice colonized with *Lactobacillus reuteri*. Pairwise comparisons of gene expression between the treatment groups were performed using DESeq2. Expression data was analyzed using the Wald test. Genes with p-values < 0.05 were defined as differentially expressed genes for each pairwise comparison. All genes differentially expressed in any pairwise comparison are included. The color scale represents the log10 (average TPM + 0.01) value. TPM: transcripts per kilobase million.

